# Characterization of Resolved, Localized, and Progressing Infections at the Maternal-Fetal Interface in a Nonhuman Primate Model

**DOI:** 10.64898/2026.09.15.751503

**Authors:** Gygeria Manuel, Celeste Coler, Jeff Munson, John Cornelius, Tyla Duell, Taeyun Kim, Andrew Vo, Austyn Orvis, Michelle Coleman, Kylee Murphy, Hong Zhao, Sidney Sun, Sophia C Chima, Vanshika Sindhu, Hazel Huang, Amanda Li, Miranda Li, Chris English, Audrey Baldessari, Briana Del Rosario, Orlando Cervantes, Raj Kapur, Lakshmi Rajagopal, Kristina M. Adams Waldorf

**Affiliations:** Department of Obstetrics and Gynecology, University of Washington, Seattle, WA, USA; Morehouse School of Medicine, Atlanta, GA, USA; School of Medicine, University of Washington, Seattle, WA, USA; Department of Psychiatry, University of Washington, Seattle, WA, USA; Center for Global Infectious Disease Research, Seattle Children’s Research Institute, Seattle, WA, USA; Department of Laboratory Medicine and Pathology, University of Washington, Seattle, WA, USA; Department of Global Health, University of Washington, Seattle, WA, USA; Department of Pediatrics, University of Washington, Seattle, WA, USA; Washington National Biomedical Research Center, University of Washington, Seattle, WA, USA; Seattle Children’s Hospital, Seattle, WA, United States

**Author notes:** ***Co-Corresponding Authors***: Kristina Adams Waldorf, MD; Box 358070, 750 Republican Ave, Seattle, WA 98109, USA; Lakshmi Rajagopal, PhD, 1916 Boren Ave, Seattle, WA 98101.

**Keywords:** placenta, chorioamnionitis, myometrium, Group B Streptococcus, nonhuman primate, preterm labor

## Abstract

**Background:** Inflammation and infection are responsible for the majority of early preterm births. Host innate immune responses at the maternal-fetal interface that contribute to sterile intra-amniotic inflammation (SIAI), bacterial clearance, or invasive infection remain poorly understood. Group B Streptococcus (*Streptococcus agalactiae*) is a leading cause of preterm birth, stillbirth, and invasive neonatal disease.

**Objective:** To determine changes in host gene expression and innate immune factors that distinguish progression from resolution of a Group B Streptococcus infection at the maternal-fetal interface in the pregnant nonhuman primate (pigtail macaque, *Macaca nemestrina*) model. We hypothesized that the host inflammatory profile that correlated with bacterial invasion and progression would be distinct from those associated with infection resolution.

**Study Design:** In a chronically catheterized pregnant pigtail macaque model, Group B Streptococcus was inoculated into the choriodecidual space. Infections were classified as progressing (invasive into the amniotic fluid), localized (bacteria persisting at the inoculation site), or resolved (minimal to no bacteria detected). Temporal and spatial RNA and protein profiles in the chorioamniotic membranes and decidua and in the myometrium were assessed by histopathology, multiplex protein assays, total RNA-sequencing, Nanostring nCounter, and spatial transcriptomics (digital spatial profiling). Differential gene expression was analyzed with limma using moderated t-tests and by gene set enrichment analysis.

**Results:** A GBS infection at the maternal-fetal interface produced three infection outcomes: progressing infections that disseminated into the amniotic fluid (AF, N=5), low-titer localized infections remaining at the inoculation site (N=6), and resolved infections (N=5), compared with saline controls (N=3). Preterm labor occurred frequently regardless of the infection outcome (40-80%). In resolved infections, SIAI occurred in the AF and lymphocytes infiltrated the myometrium. Total RNA-Seq of the chorioamniotic membranes demonstrated that 99 genes separated progressing from resolved infections, 98 of them were upregulated and dominated by IL-1 signaling and granulocyte recruitment (i.e., *IL1B*, *CXCL8*, *TLR2*, *CLEC4A*). Histone deacetylase 10 (*HDAC10*) was the only significantly downregulated gene in the progressing versus resolved contrast (p<0.05). Spatial transcriptomics localized the progressing inflammatory response to the chorion. In the myometrium, zinc and RING finger 3 (*ZNRF3*) was significantly downregulated in localized versus control and resolved versus control and was also the only differentially expressed gene. Downregulated WNT signaling and upregulated lymphocyte-mediated immunity gene sets in the myometrium were a signature of resolved infections.

**Conclusion:** A GBS infection at the maternal-fetal interface can result in a spectrum of progressing, localized and resolved outcomes. An early IL-1 and granulocyte response in the chorion layer of the chorioamniotic membranes marks bacterial dissemination, with loss of HDAC10 suggesting epigenetic regulation within the membranes with progressing infections. In the myometrium, ZNRF3 and downregulation of WNT signaling persisted even after infection resolution, reflecting an earlier infectious exposure. Unexpectedly, resolved infections were associated with SIAI and a myometrial inflammatory response, offering a possible explanation for preterm labor associated with SIAI.

## Introduction

Preterm birth is the leading cause of neonatal mortality and death globally for children under the age of five. Most early preterm births are associated with either sterile intra-amniotic inflammation (SIAI) or microbial invasion of the amniotic cavity (MIAC). Group B Streptococcus (GBS, *Streptococcus agalactiae*) is a gram-positive bacterium and a significant cause of infection and inflammation leading to preterm birth, stillbirth, and neonatal sepsis. GBS is a common colonizer of the vaginal tract; however, in pregnancy, GBS can ascend into the lower uterus, traffic into the decidua, and cross placental chorioamniotic membranes to invade the amniotic fluid (AF).^1^ An estimated 500,000 preterm births and 46,000 stillbirths annually have been linked to GBS infections worldwide.^1–6^

The relatively inaccessible maternal-fetal interface has made it challenging to understand the host innate immune responses that direct GBS bacterial clearance or aid in bacterial propagation and invasive disease. A framework for the progression or resolution of inflammation and infection has evolved over the years to incorporate several factors: (1) the type of cells responding to the infection (e.g., neutrophils); (2) the class of chemical modulators produced; (3) the degree of activation of innate immune sensors (4) the mechanism of host cell death; and (5) how dying stromal and immune cells are removed (efferocytosis).^7–9^ An invasive infection at the maternal-fetal interface is more likely to be driven by a hyperinflammatory response including cell death cascades, pyroptosis or ferroptosis^10^ Which mechanisms determine the outcome of host-pathogen responses at the maternal-fetal interface is unknown.

We have previously reported that choriodecidual GBS inoculation in a pregnant nonhuman primate (NHP) model can result in bacterial invasion of the amniotic cavity or resolve over time.^11, 12^ Here, we used the chronically catheterized NHP (pigtail macaque, *Macaca nemestrina*) model to closely examine early host innate immune responses associated with experimentally invasive and resolving GBS infections in the choriodecidual space adjacent to the uterine muscle (myometrium) and the placental chorioamniotic membranes. To elicit the temporal and spatial host responses to bacterial dissemination or resolution within the uterus and the surrounding lower uterine segment of myometrium, we inoculated GBS into our catheterized pregnant NHP model. We characterized the range of GBS infections inoculated into the choriodecidual space as either progressing (MIAC), localized (bacteria persisting primarily at the inoculation site), or resolved. In parallel, we inoculated saline into our catheterized pregnant NHP model to serve as uninfected controls. The objective was to determine the host innate immune factors and transcriptional profile that characterize the natural history of an experimental GBS infection at the maternal-fetal interface in our NHP model. We hypothesized that bacterial invasion would have a distinct host inflammatory profile from infection resolution in the placental chorioamniotic membranes and lower uterine segment myometrium.

## Materials and Methods

All animal experiments complied with the National Research Council’s Guide for the Care and Use of Laboratory Animals and the Weatherall report on NHP research. The protocol was approved by the University of Washington IACUC (Permit #4165-01; most recent approval May 7, 2025). Methods are presented in brief with full details in the supplement.

We used a chronically catheterized pregnant NHP model (pigtail macaque, *Macaca nemestrina*) to provide an experimental challenge of GBS followed by serial AF collection prior to Cesarean section (C-section) to capture early events at the maternal-fetal interface during infection. First, animals were acclimated to a jacket-tether system. Next, catheters were implanted into the maternal femoral vein, AF, and choriodecidual interface at 114-125 days gestation (term=172 days) for sampling or to monitor intrauterine pressure (Fig. 1). After an ∼10-day recovery, animals received either choriodecidual inoculation of saline (N=3) or ∼1-5 × 10⁸ CFU of GBS strain COH1*ΔcovR* (N=11) or COH1*ΔcovRΔcylE* (N=5). Preterm labor (PTL) was defined by cervical dilation with uterine activity above 10,000 mmHg·sec/hr for at least 2 hours. C-section was performed either at PTL onset or a staged study endpoint (1-3 days post-inoculation). At delivery, tissues from the maternal-fetal interface were collected and fetuses were euthanized by exsanguination and barbiturate overdose, followed by necropsy. The GBS inoculum, fluids (AF, fetal blood) and homogenized tissues (placental chorioamniotic membranes, decidua, myometrium, fetal lung, fetal brain) were cultured on TSA plates for bacterial enumeration. CAMP factor activity was assessed on sheep blood agar alongside the inoculum strain. AF was sampled at baseline (−24, −0.25 h) and post-inoculation (+0.75, +6, +12, +24 h, then every 12 h until C-section).

**Figure 1.**
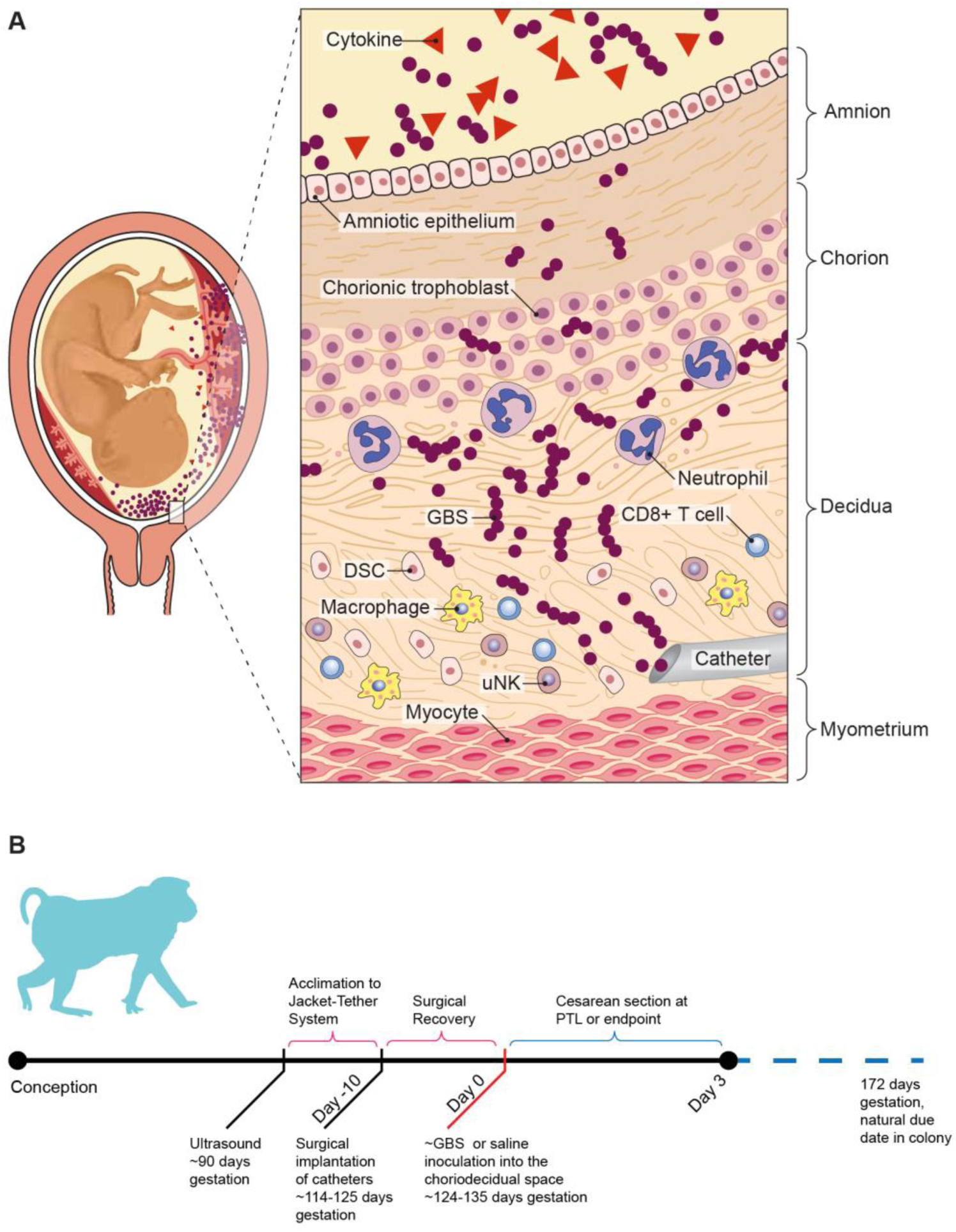
Illustration of Surgical Catheter Placement in our NHP Model and Study Design. **(A)** Schematic of surgical catheter placement via laparotomy into the maternal femoral vein, amniotic fluid, and the choriodecidual interface of the lower uterine segment at 114-125 days gestation (equivalent to human late second or early-third trimester), with the site of GBS choriodecidual inoculation indicated. **(B)** Diagram of experimental design.

Infection outcomes were defined as either: progressing (disseminated), localized to the inoculation site, or resolved. Progressing GBS infection was defined by the presence of viable bacterial counts (CFU/mL) in the AF at the end of the study, indicating spread of GBS from the inoculation site across the placental membranes, and active replication within the uterine cavity and fetus. In contrast, a resolved infection had no GBS bacteria at the inoculation site or in the AF, suggesting bacterial clearance. An intermediate outcome was defined as localized if a tissue culture was positive at the inoculation site and no viable bacteria was recovered from the AF; often, trace amounts of GBS genomic DNA in the AF was detected in this group (Table S1). During analysis, we combined localized and resolved infections into a single group, referred to as “contained” infections.

Placental and fetal histopathology was performed by a pediatric (RK) and veterinary pathologist (AB), respectively. Pro-inflammatory cytokines were measured using either Luminex multiplex assay (Millipore) or cynomolgus/rhesus ELISA kits for IFN-beta (IFN-β; PBL Assay Science, #46415). Genomic DNA was extracted (QIAamp DNA Mini Kit, Qiagen) and qPCR targeting the GBS surface immunogenic protein (SIP) gene was used to quantify trace and total bacterial DNA (Table S1). Tissues were stored in RNAlater and homogenized in TRIzol using a Precellys Evolution homogenizer. RNA was extracted with the RNeasy Mini Kit (Qiagen), quantified via Qubit 4.0, and assessed for integrity (RIN) via Agilent 4200 TapeStation.

Transcriptional profiling of the placental chorioamniotic membranes and lower uterine segment myometrium were performed using total RNA-Seq, nCounter, and spatial-Seq. Total RNA Libraries were prepared using the Roche KAPA RNA HyperPrep kit with RiboErase HMR (100 ng input, RIN>3), verified by Qubit/TapeStation, and sequenced on the NovaSeq 6000 platform. FASTQ files were aligned to the *Macaca mulatta* (rhesus macaque) reference genome (Mmul_10, Ensembl) using STAR 2.7.8a, which is a close relative to the pigtail macaque. Gene validation was performed using nCounter. Spatial-Seq was used to determine transcriptional profiles of the amnion, chorion, and decidua using formalin-fixed paraffin-embedded chorioamniotic membrane and decidua sections using morphology-guided ROI selection (Nanostring GeoMx), photocleaved oligo collection, library prep, and sequencing. Differential expression testing (total RNA-seq, nCounter, GeoMx) used limma linear models with moderated t-tests (empirical Bayes). Significance required |fold change| ≥1.5 and adjusted p<0.05 (Benjamini-Hochberg). Gene expression of HDAC10 was validated by RT-qPCR. Statistical tests used are specified in figure legends. All analyses were performed in R (v4.4.0).

## Results

### GBS infection phenotypes

We observed a spectrum of phenotypic outcomes in the first three days after an experimental GBS infection at the maternal-fetal interface of our NHP model (Fig. 1). Infections either progressed and disseminated into the AF and fetus, remained localized at the site of inoculation, or completely resolved (Table 1, Table S1-S3). The extent of bacterial dissemination did not correlate with PTL. Only 40% (2/5) in the progressing group had PTL versus 50% (3/6) in the localized, 80% (4/5) in the resolved group, and 0% (0/3) in the uninfected control group (Table S3). Viable, replicating GBS in the AF was only detected in the progressing group, although trace genomic DNA from GBS was detected in the localized group. The progressing group was the only group where fetuses were infected with GBS.

**Table 1.** GBS infection groups and associated adverse pregnancy outcomes.

| Group | PTL Frequency | GBS Detection by Swab of the Choriodecidual Inoculation Site | Mean GBS in AF by Culture (CFU/mL) | Mean GBS SIP DNA in AF by qPCR (GE/mL) | Mean GBS in Fetal Lung by Lung Culture (CFU/mL) |
| --- | --- | --- | --- | --- | --- |
| Control (N=3) | 0% | Negative | 0 | 0 | 0 |
| Resolved (N=5) | 80% | Negative | 0 | 0 | 0 |
| Localized (N=6) | 50% | Usually positive | 0 | $6.1 \times 10^2$ | 0 |
| Progressing (N=5) | 40% | Positive | $2.9 \times 10^7$ | $2.2 \times 10^7$ | $9.2 \times 10^4$ |

This table shows the defining features of a progressing, localized, and resolved GBS infection based on average AF culture, swab of the inoculation site, and average GBS quantitative PCR (qPCR) in the AF. It also shows the rate of associated preterm labor. Abbreviations: AF, amniotic fluid; CFU, colony-forming units; GE, genomic equivalents.

Histopathology review of the placental chorioamniotic membranes and decidua at the inoculation site revealed extensive neutrophilic infiltration of decidua and chorion in the progressing group versus minimal or mild infiltration in localized and minimal in the resolved group (Fig. 2). In contrast, there was lymphocytic infiltration in the myometrium of the resolved GBS infections adjacent to the inoculation site, which was not observed in the progressing infections. The localized group appeared either similar to the resolved or had mild lymphocytic infiltration.

**Figure 2.**
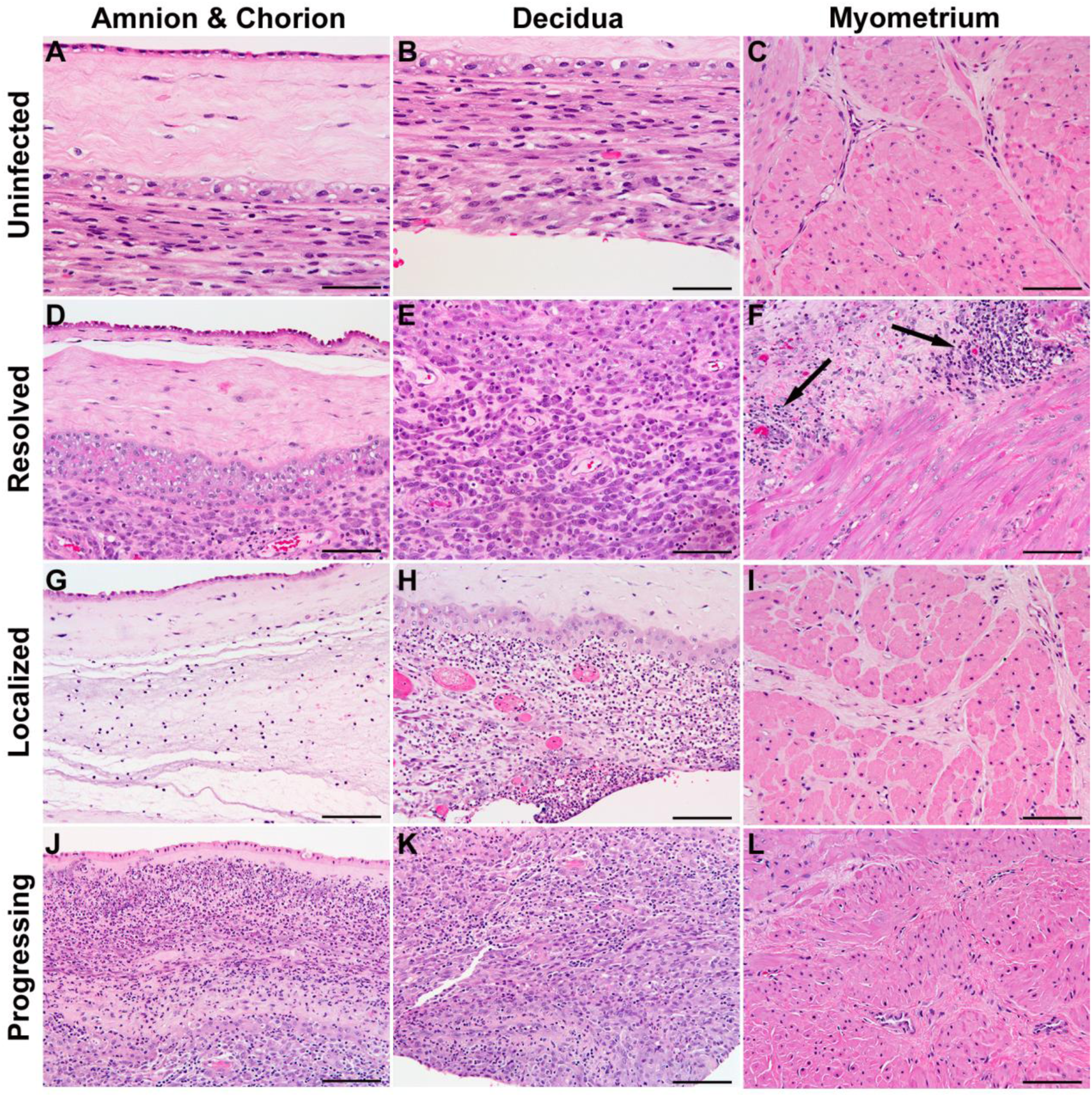
Histopathology of the Placental Chorioamniotic Membranes, Decidua, and Myometrium in the Nonhuman Primate Model of GBS Infections. Representative histological findings in the chorioamniotic membranes/ decidua and myometrium from uninfected controls (A-C), and GBS infections determined to have either resolved (D-F), remained localized at the choriodecidual inoculation site (G-I), or progressed (J-L) to invade the amniotic cavity and fetus. Images show the amnion and chorion (A, D, G, J), the decidua (B, E, H, K), and the myometrium (C, F, I, L). Dense aggregates of lymphocytes (arrows, F) are present in the connective tissue between and overlying smooth muscle cells in a resolved animal. Scale bars: 100 µm.

To determine the cytokine profile of tissues at the maternal-fetal interface by infection group, we quantified cytokines in the AF and chorioamniotic membranes and lower uterine segment of the myometrium at the study endpoint (Fig. 3, Fig. S1-S3). The most striking difference in pro-inflammatory cytokines was the significant elevation in IL-1 beta (IL-1β), and TNF across the AF, placental chorioamniotic membranes, and lower uterine segment myometrium between the progressing and uninfected control group at 24 hours post-inoculation (p<0.05, all contrasts). IL-6 was also significantly elevated in the AF between progressing and uninfected groups at this timepoint (p<0.05). Similar cytokines were significantly elevated in the progressing versus the resolved group: IL-1β (AF, membranes), TNF (AF), and IL-6 (membranes, all p<0.05). Other cytokines exhibited trends toward increased expression during progressing infection but were not statistically significant (Fig. S1-S3). These findings indicate a greater pro-inflammatory response associated with GBS infection progression that is most evident within the AF and chorioamniotic membranes but also present in the adjacent myometrium. IL-1-driven inflammatory pathways were a key feature of progressing/disseminated GBS infections.

**Figure 3.**
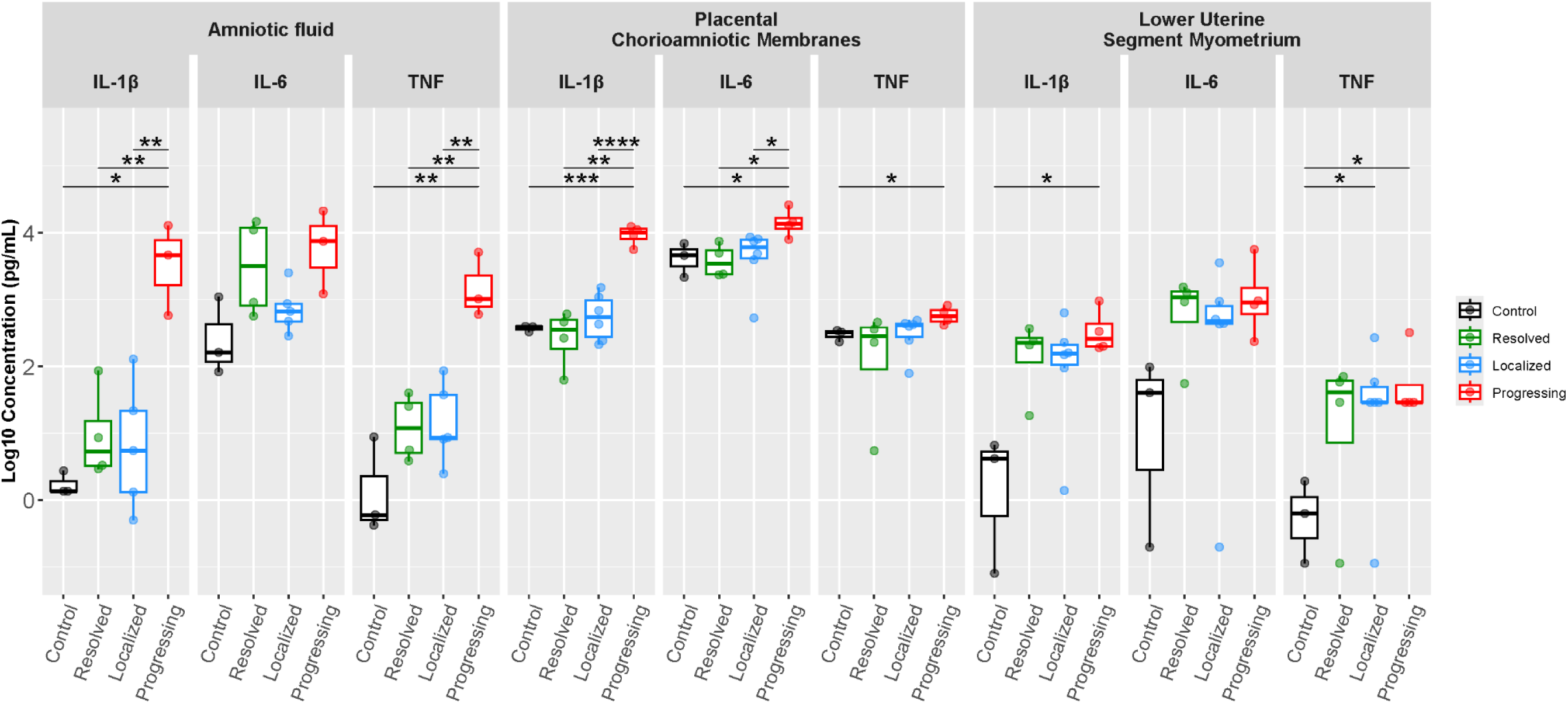
Pro-Inflammatory Cytokines in the Amniotic Fluid and Placental Chorioamniotic Membranes at the GBS Inoculation Site. The box plots show pro-inflammatory cytokine concentrations of IL-1β, IL-6, and TNF from a 14-plex panel in amniotic fluid at 24-hours post inoculation or in the placental chorioamniotic membranes and the lower uterine segment myometrium at the study endpoint (preterm labor or 3 days post-inoculation. The x-axis reflects the experimental groups, and the colors indicate uninfected controls (black), resolved (green), localized (blue) and progressing (red). The y-axis is the log10 of the cytokine concentration (pg/mL). Student’s t-test was performed on log10 transformed values. A single star reflects a p-value <0.05, and double stars reflect a p-value <0.01.

### Bulk RNA-Seq Transcriptional Profiles of Chorioamniotic Membranes

To determine whether infection phenotypes reflected differences in transcriptional profiling, we performed total RNA-Seq on the placental chorioamniotic membranes, noting that LOC1 did not contribute data due to a missing sample (Fig. 4, Fig. S4-S10). There were 3,199 differentially expressed genes (DEGs; fold change > |1.5|) in the progressing versus control group (Fig. 4A, S5). Granulocyte recruitment and IL-1 signaling dominated the profile (e.g., matrix metalloproteinase 8, *MMP8*: L2FC=10.3, p=0.04; *IL1B*: L2FC=6.7, p=0.001). Localized infection produced an intermediate response, with 1,649 DEGs (Fig. 4A, S6). Granulocyte and IL-1 signaling transcripts were increased, but not significant (*MMP8*: L2FC=1.5, p=0.86; *IL1B*: L2FC=1.0, p=0.51). There were 2,997 DEGs in the resolved group with even lower expression of key granulocyte and IL-1 transcripts (*MMP8*: L2FC=1.4, p=0.83; *IL1B*: L2FC=0.7, p*=*0.59; Fig. 4A, Fig. S7). A similar pattern of greater DEG expression in the progressing versus control group was apparent for genes linked to microbial sensing, the NF-kappa B (NF-κB) pathway, and matrix degradation (Fig. 4A).

**Figure 4.**
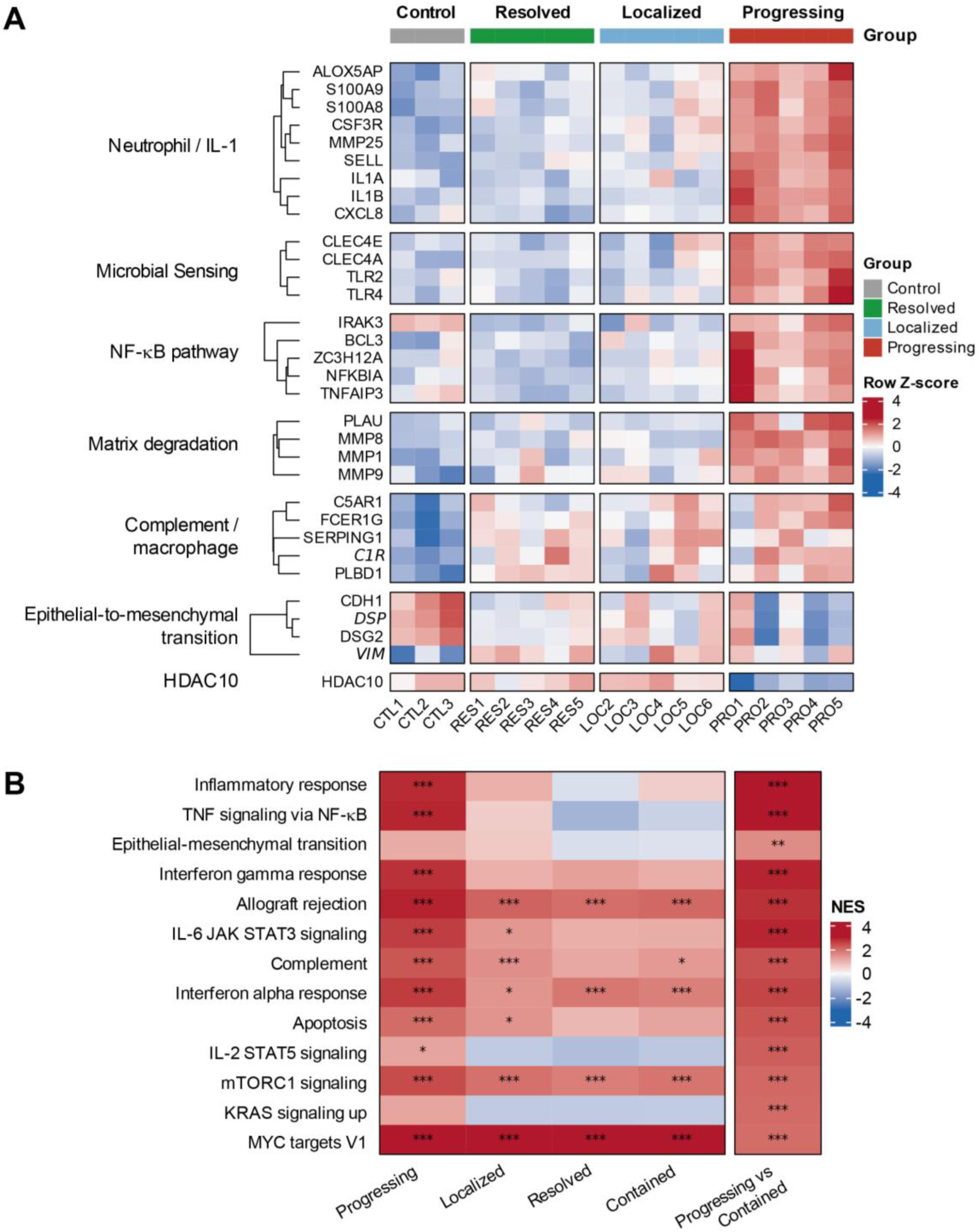
Total RNA-Seq Analysis of the Placental Chorioamniotic Membranes by Infection Group. This figure illustrates the gene-level (A) and gene set-level analysis (B) of differential expression between the progressing versus control, localized versus control, resolved versus control, and progressing versus contained (localized + resolved). Within both panels, red indicates an upregulated gene or gene set and blue corresponds to a downregulated gene or gene set. The white color indicates no differential regulation between the two groups. In panel A, a colored band at the top of the heatmap indicates the experimental group: controls (gray), resolved (green), localized (blue), and progressing (red). Each row represents an individual gene, and each column represents the expression level for an individual animal’s chorioamniotic membrane sample. Genes were grouped into distinct functions or pathways listed on the left. The color of each box of the heatmap represents the Z-score calculated by averaging the expression level across all the samples in a single row. In panel B, a GSEA was performed using pathways from the MSigDB Hallmark gene sets, indicated on the left. Each column represents the pathway expression level of an infected group (listed on the bottom) compared to controls, except for the last column which compares progressing versus contained groups. The color of each box of the heatmap represents the normalized enrichment score (NES) and the adjusted p-value of each comparison is indicated by the stars (1 star: p<0.05, 2 stars: p<0.01, 3 stars: p<0.001). Note that all animals except one (LOC1) contributed total RNA-Seq data.

Genes associated with complement and macrophage activation were more similarly upregulated in all experimental groups (Fig. 4A). In contrast, there was a loss of DEGs maintaining epithelial integrity and a concomitant gain of vimentin across all infected groups, consistent with epithelial-to-mesenchymal (EMT) transition, which might weaken membrane strength (i.e., progressing versus control: E-cadherin, *CDH1*, L2FC=-3.7, p=0.016; *VIM*: L2FC=1.1, p=0.05). Histone deacetylase 10 (*HDAC10*) had the greatest reduction in expression for progressing versus resolved (L2FC=-0.98, p=0.02), and this result was validated by RT-qPCR (Fig. 4A, S8, S11). HDAC10 was also significantly downregulated in progressing versus localized groups (L2FC=-1.08, p=0.02), although this was not significant by RT-qPCR. (Fig. 4A, S9, S11).

Direct comparison of the progressing group to either the localized or resolved groups revealed a narrow, IL-1 signaling and granulocyte recruitment signature. There were 99 DEGs in the progressing versus resolved contrast, 98 of which were upregulated and dominated by IL-1 signaling (Fig. S8; *IL1B*: L2FC=6.0, p<0.001). Only 40 DEGs differentiated progressing from localized infections, of which 39 were upregulated and from the same IL-1 and granulocyte recruitment pathways (*IL1B*: L2FC=5.8, p<0.001; Fig. S9). Comparing the resolved and localized groups directly revealed no significant DEGs (Fig. S10). Therefore, the localized and resolved groups were combined into a new ‘contained’ group for Gene Set Enrichment Analysis (GSEA, Fig. 4B). These analyses confirmed a significant enrichment of the neutrophil, IL-1 signaling, toll-like receptor, and pathogen recognition receptor signaling pathways in the progressing group versus controls or the contained infections (Fig. 4B, all p<0.05). Macrophage activation, complement, phagocytosis, and EMT gene sets were also significantly enriched in the progressing versus contained group (p<0.05).

### Spatial GeoMx Transcriptional Profiles of Amnion, Chorion, and Decidua

Next, we investigated spatial gene expression differences to determine sites within the chorioamniotic membranes and decidua that were responsible for the spectrum of differential gene expression observed across the group. We used the GeoMx platform to define regions of interest (ROIs) and profile gene expression within the amnion, chorion, and decidua (Fig. 5, 5A-B). Transcripts within the amnion were sparse and insufficient for analysis. Comparting the progressing versus contained groups, we asked whether genes expressed in the chorion or decidua were responsible for the signature observed in the bulk RNA-Seq analysis. The neutrophilic/IL-1 signature and inflammatory response observed in the progressing versus contained group was derived predominantly from the chorion (Fig. 5C-D) with a lesser contribution of the decidua (Fig. 5E-F). Other gene sets in the chorion, which were also significantly upregulated for this contrast included complement, allograft rejection, and IL6 JAK/STAT3 signaling (Fig. 5D). In the decidua, inflammatory gene sets were upregulated but to a lesser degree than the chorion (Fig. 5F).

**Figure 5.**
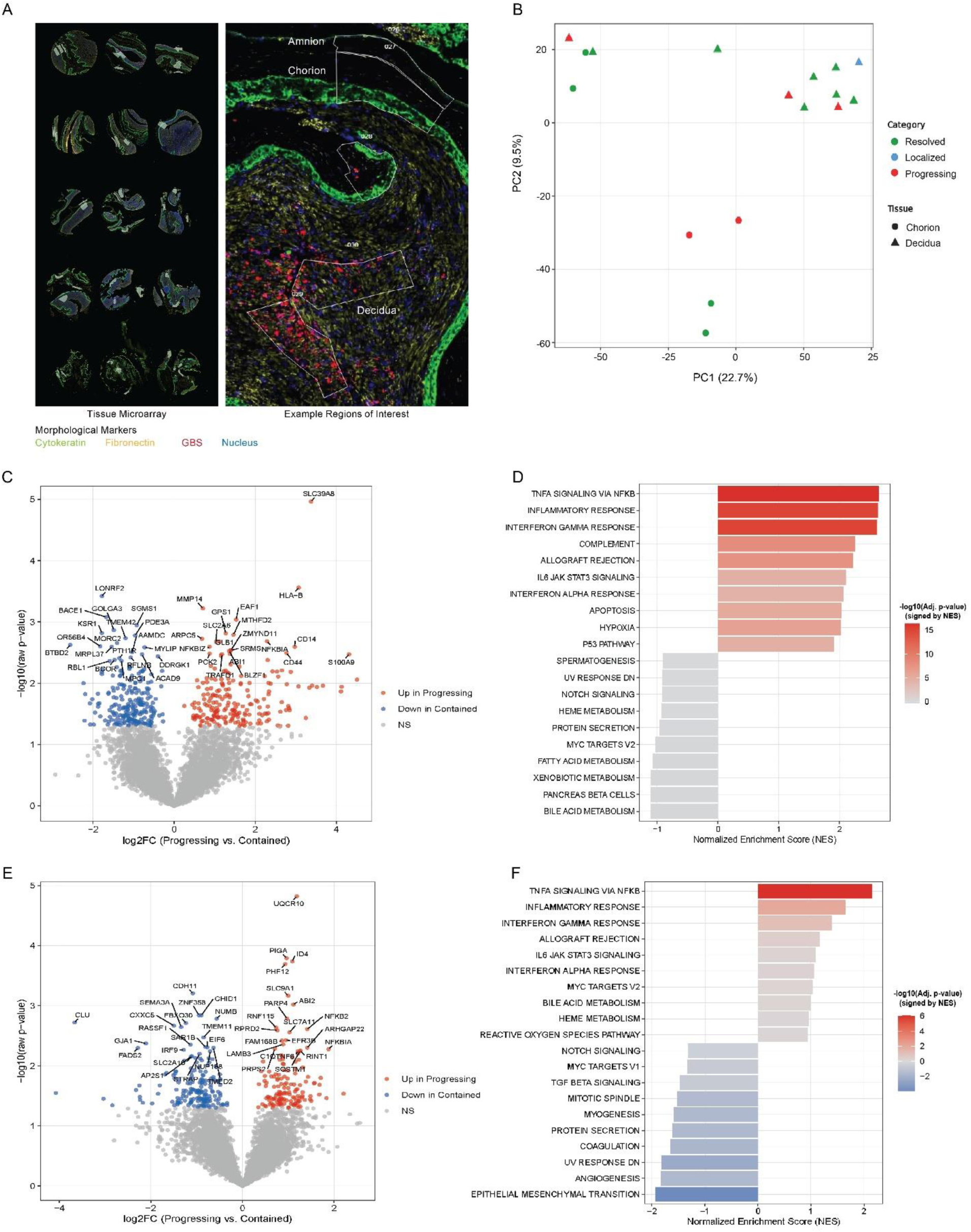
Spatial Transcriptomics of the Amnion, Chorion, and Decidua Between Progressing and Contained GBS Infections at the Placental Chorioamniotic Membrane Inoculation Site. This figure depicts the GeoMx spatial transcriptomics analysis of amnion, chorion, and decidua samples in GBS-infected animals separated into progressing and contained groups. A) Regions of interest defined by morphological markers (cytokeratin: Pan-CK, green; fibroblast activation protein: FAP, yellow; nucleus: DAPI, blue; GBS: anti-GBS, red). B) Principal component analysis (PCA) of DESeq2 normalized counts of spatial transcriptomics performed on chorion (triangles) and decidua (squares) in contained (blue) and progressing (red) groups. Volcano plots for chorion (C) and decidua (E) comparing progressing vs contained groups with log2 fold-change on the x-axis and –log10 raw p-value on the y-axis. Red dots represent genes that were significantly upregulated in the progressing versus contained group, while blue dots represent significantly downregulated genes (raw p<0.05, fold change > |1.5|). A GSEA bar plot showing the top 10 enriched pathways in the chorion (D) and decidua (F). Pathways are ordered by the normalized enrichment score (NES), with red bars indicating pathways that are upregulated in progressing versus contained, and blue bars indicating pathways that are downregulated. All pathways were analyzed based on the MSigDB Hallmark gene sets. The intensity of the red or blue color represents the magnitude of the –log10 adjusted p-value.

### Bulk RNA-Seq Transcriptional Profiles of Lower Uterine Segment Myometrium

Next, we analyzed total RNA-Seq profiles of the lower uterine segment myometrium adjacent to the inoculation site to determine if transcriptional responses could provide insight on how the spectrum of GBS infections influences the risk of PTL (Fig. 6, S12). Zinc and RING finger 3 (*ZNRF3*) was significantly downregulated in localized versus control and resolved versus control comparisons and was the only differentially expressed gene. At the individual gene level, Zinc and RING finger 3 (*ZNRF3*) was the only transcript meeting significance, and it was downregulated in each infected group relative to controls. The GSEA revealed a significant downregulation of multiple gene sets related to WNT signaling pathways; notably, *ZNRF3* encodes a transmembrane E3 ubiquitin ligase, which negatively regulates Wnt/β-catenin signaling. Enrichment of gene pathways related to lymphocyte-mediated immunity was also observed, particularly in the resolved versus control contrast. Downregulation of *ZNRF3* and WNT signaling across all infections groups versus controls indicates it is a marker of GBS exposure that persists after infection resolution, while enrichment of adaptive immunity in resolved versus control matches the histopathologic findings of leukocyte infiltration in resolved animals.

**Figure 6.**
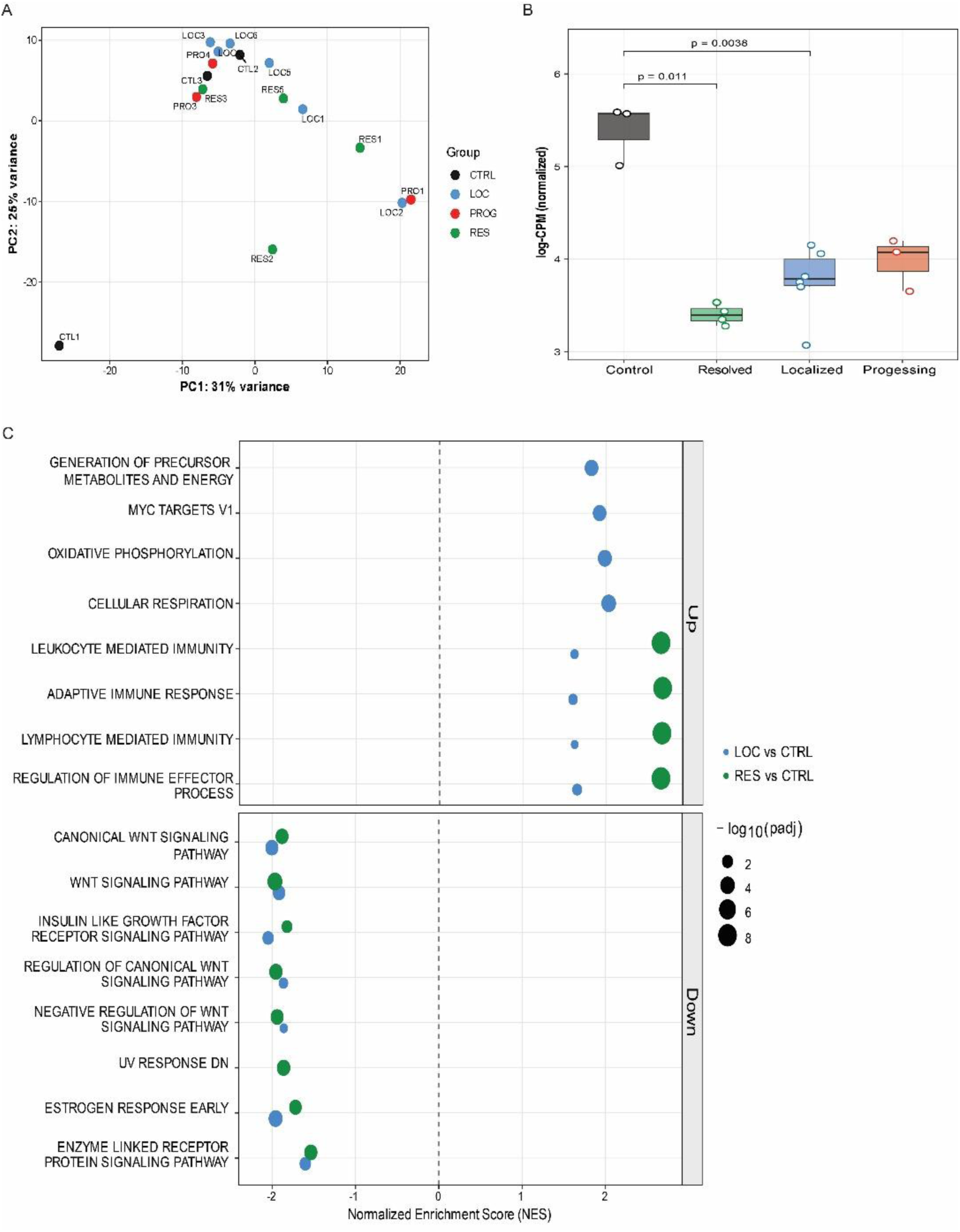
Total RNA-Seq Analysis of GBS Infections and Uninfected Controls in the Myometrium Lower Uterine Segment Adjacent to the Inoculation Site. This figure presents total RNA-Seq results from the lower uterine segment myometrium. A) A Principal Component Analysis plot shows clusters of animals by experimental group (black: control; green: resolved; blue, localized; red, progressing). B) A box plot demonstrates that ZNRF3 gene counts were significantly less in the myometrium of the lower uterine segment in both localized versus control and resolved versus control (all, p<0.05). C) GSEA of a localized versus control (blue) and resolved versus control (green) shows significant upregulation of the adaptive immune response in resolved infections and downregulation of WNT signaling pathways for both contrasts.

## Comment

### Principal Findings

Our results demonstrate that GBS infections at the maternal-fetal interface can progress and disseminate into the AF or resolve, which leaves a residual, sterile inflammatory response in the AF and myometrium. IL-1 signaling and granulocyte recruitment developed rapidly during progressing infections in the chorioamniotic membranes and were established by 24 hours post-inoculation. Resolved and localized infections were associated with similar transcriptional profiles suggesting that localized infections that had not disseminated in the first few days were on a path towards resolution. Spatial transcriptomics localized the inflammatory response primarily to the chorion. The significance of the leukocytic infiltrate in the myometrium of resolved infections is unclear but may have contributed to bacterial clearance and the high PTL rate in this group.

### Comparison with Existing Literature

Understanding the natural history of infections at the maternal-fetal interface within days after onset is not possible in human pregnancy. Murine models of pregnancy do not recapitulate key features of human pregnancy (i.e., placentation) or human immunology, which would make it nearly impossible to perform a similar study and translate the findings.^13–16^ Although murine models of preterm birth exist, infections are supraphysiologic and often cause stillbirth, which makes it impossible to separate infection-driven myometrial activation from biological events associated with stillbirth and placental failure.

As microbe-host interactions at the maternal-fetal interface have not previously been studied in this manner, it has been assumed that bacterial infections overwhelmed placental defenses to necessarily infect the AF and fetus. Although we have previously shown that GBS infections at the maternal-fetal interface can resolve^11^, the host response associated with the spectrum of GBS infection outcomes was unknown. While the progressing group recapitulates an early preterm birth phenotype with MIAC, the data linking a resolved infection to SIAI and preterm labor is novel. These data suggest that SIAI, a common finding in women with PTL in the late second and early third trimesters, may have microbial origins.^17–21^ Immune cells that are more numerous or more functionally competent in late gestation may favor infection resolution and a sterile inflammatory response signature, which could explain why SIAI predominates in late gestation, while MIAC is more common earlier.^19, 22^ In this context, post-infectious SIAI represents a snapshot in time, which follows an infection but retains a microbial signature that outlasts viable bacteria. If SIAI has a microbial origin, this would dramatically increase the number of PTL cases associated with an infectious cause.

### Strengths and Limitations

The strength of this study is in the ability to follow the natural history of a controlled infection at the maternal-fetal interface in a NHP model that is most similar to human pregnancy.^13^ Delivery within the first 3 days after choriodecidual inoculation, and natural variation in infection outcome, allowed the investigation of transcriptional responses in the chorioamniotic membranes and myometrium associated with progressing versus contained infections for the first time. We also recognize the study’s limitations. Defining the signaling pathways that trigger or contain bacterial dissemination versus those that are induced by the outcome itself may not be possible. A conservative interpretation of our results is that the transcriptional signatures identified reflect a downstream event of bacterial dissemination or containment, rather than a cause. Even with this important limitation, the data brings us closer to understanding how host-pathogen interactions at the maternal-fetal interface might manifest for a pregnancy. The knowledge that a resolved infection can trigger PTL leaving behind an adaptive immune response within the myometrium and a sterile inflammatory response in the AF, is a major advance in understanding the origins of late preterm birth, which is often characterized by SIAI. Another limitation is the use of two different GBS strains; however, both strains contributed to progressing and contained infections.

### Conclusions

There was a spectrum of outcomes to a controlled GBS infection at the maternal-fetal interface resulting in progressing, localized, and resolved infections. Resolved infections were associated with SIAI and leukocytic infiltrates in the myometrium, whereas progressing infections were linked to an inflammatory IL-1 response. Unexpectedly, PTL occurred in both groups and was more frequent after resolution than progression (80% versus 40%). An IL-1 signature was already established in progressing infections by 24 hours post-inoculation, indicating a rapid onset after inoculation. Additional studies are needed to test preterm birth therapeutics to support infection resolution while suppressing inflammation that can trigger PTL. Understanding the spectrum of infections at the maternal-fetal interface is crucial to reducing the risk of preterm birth.

## Acknowledgments

We thank Riley Raker for assistance with graphical design.

## Author Contributions (CRediT)

GM: Methodology, Investigation, Formal analysis, Writing – original draft, Writing – review and editing. CC: Investigation, Formal analysis, Writing – original draft, Writing – review and editing. JM: Methodology, Investigation, Data curation, Visualization, Resources, Writing – review and editing. JC: Methodology, Investigation, Data curation, Visualization, Resources, Writing – review and editing. TD: Investigation, Data curation, Writing – review and editing. TK: Data curation, Writing – review and editing. AV: Methodology, Investigation, Data curation, Writing – review and editing. AO: Investigation, Writing – review and editing. MC: Investigation, Writing – review and editing. KM: Investigation, Validation, Writing – review and editing. HZ: Investigation, Validation, Writing – review and editing. SS: Investigation, Visualization, Writing – review and editing. SC: Investigation, Visualization, Writing – review and editing. HH: Investigation, Writing – review and editing. AL: Investigation, Writing – review and editing. ML: Investigation, Writing – review and editing. CE: Investigation, Writing – review and editing. AB: Investigation, Writing – review and editing. BDR: Investigation, Data Curation, Writing – review and editing. OC: Investigation, Validation, Writing – review and editing. RK: Investigation, Visualization, Writing – review and editing. LR: Conceptualization, Funding acquisition, Supervision, Resources, Writing –review and editing. KMAW: Conceptualization, Funding acquisition, Supervision, Resources, Writing – original draft, Writing –review and editing

## Data Availability

Single-cell RNA sequencing data generated in this study have been deposited in the NCBI Gene Expression Omnibus (GEO) under accession number GSE347154. All remaining data supporting the findings of this study, including source data for figures are available in the Dryad Digital Repository at https://doi.org/10.5061/dryad.xwdbrv1s9. Any additional information required to reanalyze the data reported in this paper is available from the corresponding author on request.

## Declaration of Generative AI and AI-Assisted Technologies in the Manuscript Preparation Process

During the preparation of this work the authors used Claude (Anthropic) to edit and condense the abstract and portions of the Results text for clarity and length. After using this tool, the authors reviewed and extensively edited the content. They take full responsibility for the content of the published article. No generative AI tools were used in analyzing the data or producing the figures.

