## Supplemental Information for "Characterization of Resolved, Localized, and Progressing Infections at the Maternal-Fetal Interface in a Nonhuman Primate Model"

### Table of Contents

MATERIALS AND METHODS----------1

TABLE S1. GBS SIP RT-QPCR PRIMERS AND PROBES ----------7

TABLE S2. DEMOGRAPHICS AND EXPERIMENTAL CHARACTERISTICS ----------8

TABLE S3. METHOD OF GBS DETECTION AND PRETERM LABOR STATUS IN THE CONTROL AND INFECTION GROUPS ----------10

FIGURE S1. CYTOKINES IN AMNIOTIC FLUID. ----------11

FIGURE S2. CYTOKINES IN THE PLACENTAL CHORIOAMNIOTIC MEMBRANES. ----------12

FIGURE S3. CYTOKINE CONCENTRATIONS IN THE LOWER UTERINE SEGMENT MYOMETRIUM. ----------13

FIGURE S4. PCA OF TOTAL-RNA-SEQ FROM THE PLACENTAL CHORIOAMNIOTIC MEMBRANES ----------14

FIGURE S5. TOTAL RNA-SEQ ANALYSIS OF PROGRESSING GBS INFECTIONS VERSUS CONTROL IN THE PLACENTAL CHORIOAMNIOTIC MEMBRANES AT THE INOCULATION SITE ----------15

FIGURE S6. TOTAL RNA-SEQ ANALYSIS OF LOCALIZED GBS INFECTIONS VERSUS CONTROL IN THE CHORIOAMNIOTIC MEMBRANES AT THE INOCULATION SITE----------17

FIGURE S7. TOTAL RNA-SEQ ANALYSIS OF RESOLVED GBS INFECTIONS VERSUS CONTROL IN THE PLACENTAL CHORIOAMNIOTIC MEMBRANES AT THE INOCULATION SITE----------19

FIGURE S8. TOTAL RNA-SEQ ANALYSIS OF PROGRESSING GBS INFECTIONS VERSUS RESOLVED IN THE PLACENTAL CHORIOAMNIOTIC MEMBRANES AT THE INOCULATION SITE ----------21

FIGURE S9. TOTAL RNA-SEQ ANALYSIS OF PROGRESSING GBS INFECTIONS VERSUS LOCALIZED IN THE CHORIOAMNIOTIC MEMBRANES AT THE INOCULATION SITE----------23

FIGURE S10. TOTAL RNA-SEQ ANALYSIS OF LOCALIZED GBS INFECTIONS VERSUS RESOLVED IN THE CHORIOAMNIOTIC MEMBRANES AT THE INOCULATION SITE----------25

FIGURE S11. GENE EXPRESSION OF HDAC10 BY RT-QPCR IN CHORIOAMNIOTIC MEMBRANES AT THE INOCULATION SITE----------26

PAGEREF _TOC240200942 \H FIGURE S12. TOTAL RNA-SEQ VOLCANO PLOTS OF GBS INFECTIONS AND UNINFECTED CONTROLS IN THE MYOMETRIUM LOWER UTERINE SEGMENT ADJACENT TO THE INOCULATION SITE ----------27

### Materials and Methods

#### *Research Animals and Ethics Approval*

Animal experiments were conducted in full compliance with the guidelines outlined in the Guide for the Care and Use of Laboratory Animals by the National Research Council and “The Use of Non-Human Primates in Research” in the Weatherall report. The study protocol was approved by the University of Washington Institutional Animal Care and Use Committee under Permit Number 4165-01, with the most recent approval on May 7, 2025. Surgeries were performed under general anesthesia, and every precaution was taken to minimize animal distress. All experimental animals were pregnant pigtail macaques (*Macaca nemestrina*) (Table S2). Both female animals (dams) and their fetuses were studied; the model is inherently female, and fetal sex was recorded.

#### *Pregnant Nonhuman Primate Model of Preterm Labor*

The chronically catheterized pregnant nonhuman primate (NHP) model of preterm labor (PTL) has been used in our laboratory for over two decades. Pregnant NHPs underwent surgical implantation of catheters via laparotomy into the maternal femoral vein, amniotic fluid, and choriodecidual interface in the lower uterine segment between 114-125 days gestation (term=172; ~late second or early third trimester in humans). The NHPs received either a choriodecidual inoculation of the GBS strain COH1ΔcovR of 1–5 × 10^8 colony-forming units (CFU; n=11) or GBS strain COH1ΔcovRΔcylE of 1–5 × 10^8 CFU (n=5). Experimental controls were also catheterized and received saline inoculations (n=3).

#### *Preterm Labor and Cesarean Section*

Preterm labor was defined as cervical change with increased uterine activity (>10,000 mmHg·sec/hr sustained over at least 2 hours). Cesarean sections were performed at either (1) preterm labor, (2) the designated endpoint of 1 or 3 days post-inoculation, or (3) 1 day after saline inoculation. Fetuses were euthanized by exsanguination followed by barbiturate overdose prior to fetal necropsy. Tissue collection took place at the endpoint, followed by complete gross and histopathologic examination. A 1- or 3-day endpoint was designed to investigate the earliest events and effects of GBS infection on placental tissues, fetal injury, and infection/inflammation-mediated preterm birth pathways. One-day control NHP experiments with saline choriodecidual inoculation were performed to more closely match the study design of the GBS 1-day inoculations. There was no inflammation and minimal signs of preterm labor in the one-day controls.

Intra-amniotic pressure was continuously recorded using an implanted amniotic fluid pressure catheter (SPR-524, ADInstruments, Colorado Springs, Colorado, USA) and digitized with a Powerlab System (ADInstruments). The amniotic fluid pressure signals were analyzed using custom software to eliminate noise due to respiration or position changes. The area under each contraction (mmHg·sec/hr) was summed for each hour, allowing calculation of the hourly contraction area, a measure of uterine activity. We then calculated the mean hourly contraction area over a 24-hour period. Cervical dilation and length were recorded at baseline and delivery.

#### *Bacterial Growth and Confirmation of Infection*

The GBS isogenic strains COH1ΔcovR and COH1ΔcovRΔcylE were derived from wild-type GBS COH-1, an ST-17 clone belonging to capsular serotype III, recovered from a neonate with meningitis. Single colonies of minimally passaged GBS from blood agar plates were grown in tryptic soy broth or tryptic soy agar at 37°C in 5% $CO_2$ overnight. The GBS strains were grown to mid-log phase ($OD_{600}$ = 0.5) and prepared to a concentration of approximately 1–5 × 10^8 CFU in 1 mL PBS before inoculation into the choriodecidual space.

Aliquots of the prepared inoculum were plated on tryptic soy agar (TSA) to confirm strains and assess the precise inoculation amount. On TSA, COH1ΔcovR appears orange and COH1ΔcovRΔcylE appears white. Both strains were spectinomycin-resistant and thus were subcultured on selective medium (i.e., TSA containing spectinomycin). In addition to the inoculum, amniotic fluid, fetal blood, homogenized placental (chorioamniotic membranes and decidua) and fetal tissues (lung, brain, fetal blood) were plated on TSA plates and enumerated. The level of CAMP factor activity was measured on sheep blood agar plates with the inoculum strain included in parallel.

#### *Sample Collection and Processing*

Amniotic fluid (AF) was collected throughout the experimental study. AF was sampled at baseline (−24 and −0.25 h) and post-GBS inoculation (+0.75, +6, +12, +24 h and then every 12 h until Cesarean section for fetal necropsy). Viable GBS bacteria were determined by AF serial dilutions plated on TSA, incubated overnight at 37°C, 5% $CO_2$, and enumerated. Additional aliquots were flash-frozen at −80°C. For detection of low GBS amounts in AF, qPCR (DNA quantification) was performed on endpoint AF samples. For quantification of cytokine and chemokine concentrations, AF samples were collected in EDTA. Immediately after sample collection, samples were centrifuged for 5 min at 48 × g. The supernatants were frozen and stored at −80°C. Similar to AF, viable GBS in fetal blood were determined by serial dilutions plated on TSA, incubated overnight at 37°C, 5% $CO_2$, and enumerated.

In all experiments, fetal and placental tissues (chorioamniotic membranes and decidua at the inoculation site and lower uterine segment of myometrium) were either flash-frozen and stored at −80°C, preserved in RNAlater and then frozen at −80°C, or placed in formalin-fixed paraffin-embedded molds. Flash-frozen samples were used for the quantification of cytokine and chemokine concentrations. Samples stored in RNAlater were used for gene expression analysis, total RNA sequencing, and nCounter, as well as the detection of bacterial genomic DNA. Formalin-fixed embedded tissues were used for Nanostring GeoMx Digital Spatial Profiling Whole Transcriptome Analysis.

#### *Cytokine Quantitation*

Immune mediators at the tissue level were revealed using multiplex and simplex immunoassays validated for primate species. Cytokine and chemokine concentrations were determined by Luminex multiplex technology using commercially available non-human primate cytokine kits (Millipore, Billerica, MA). Interferon-beta (IFN-β) was determined using a commercially available cynomolgus/rhesus ELISA kits (PBL Assay Science, cat #46415).

#### *GBS DNA qPCR in Amniotic Fluid, Myometrium, and Chorioamniotic Membranes/Decidua*

GBS qPCR was used to detect both trace and total amounts of bacterial genomic DNA in AF and tissue samples. Primers and probes were designed using Integrated DNA Technologies (San Diego, CA) based on validated experiments on the surface immunogenic protein (SIP) found in GBS (Table S1). SIP is conserved in all GBS strains, including COH1ΔcovR and COH1ΔcovRΔcylE. TaqMan Fast Advanced Master Mix (Fisher Scientific, cat #444964) was used for amplification. qPCR was performed on the

QuantStudio 3 machine (Thermo Fisher Scientific, cat #A28132) according to the manufacturer's instructions. Samples were run in triplicate. Positive results were defined as any sample with an average Ct value of 38 cycles or less (Table S3).

Genomic DNA from amniotic fluid and placenta tissues was extracted using the QIAamp DNA Mini Kit (Qiagen, cat #51306). For amniotic fluid, equal volumes of sample and AL buffer were mixed. Proteinase K was added at 1:10 to the volume of AL buffer (e.g., 20 µL Proteinase K to 200 µL AL buffer). Samples were incubated at 56°C for 10 minutes. DNA was eluted in 50 µL Qiagen elution buffer. DNA was extracted from myometrium homogenates prepared in TRIzol. Tissue samples were mixed with 500 µL of Back Extraction Buffer (BEB; 4 M guanidine thiocyanate, 50 mM sodium citrate, NaCl, and 1 M Tris base, prepared in ddH2O) and centrifuged at 12,000 × g for 30 minutes. The aqueous upper phase was resuspended with 400 µL of ice-cold isopropanol to precipitate DNA. Samples underwent two wash steps that included centrifugation (12,000 × g for 15 minutes at 4°C) and resuspension with ethanol. The final DNA pellet was dissolved in 30 µL of Qiagen elution buffer. Amniotic fluid and myometrium DNA were quantified using a Nanodrop spectrometer.

#### *Definition of GBS Groups in the Pigtail Macaque*

GBS-infected NHPs were categorized into three groups based on bacterial enumeration (viable GBS) and GBS DNA (total GBS) counts in AF (Table S3). Animals with a progressing GBS infection had viable counts (minimal concentration of $5 \times 10^5$ CFU/mL) and at minimum $1 \times 10^6$ genomic DNA equivalents per mL (GE/mL; total GBS DNA amount) in the AF. A localized GBS infection was defined by positive culture results at the inoculation site in the choriodecidual space or the chorioamniotic membranes but no viable counts in the AF; a trace amount of GBS DNA in the AF by qPCR (~10-1,000 GE/mL) was determined. This was deemed to represent an infection intermediate between progressing and resolved. In contrast, a resolved GBS infection was defined by no viable counts of GBS in the choriodecidual space, chorioamniotic membranes, or AF by bacterial plating or qPCR.

#### *RNA Isolation*

High-quality RNA was required to investigate transcriptome changes using the total RNA-sequencing and nCounter platforms. Tissue samples stored in RNAlater were combined with TRIzol (1 mL) and homogenized by the Precellys Evolution machine (Bertin Technologies, cat #K002198-PEVO0-A.0). RNA was extracted from tissue samples using the RNeasy Mini Kit (Qiagen, cat #74104). The concentration of RNA was determined by the Qubit 4.0 fluorometer (Thermo Fisher, cat #Q33238), and the integrity of RNA (RIN) was measured by the Agilent 4200 TapeStation (Agilent, cat #G2991BA).

#### *Total RNA Sequencing and Analysis*

Total RNA-seq was performed to capture a broad and unbiased view of transcriptional changes in the placenta and myometrium. The Roche KAPA RNA HyperPrep kit with RiboErase HMR (cat #8098131702) was used for library preparation with an RNA input of 100 ng and RIN >3. RiboErase HMR was used to reduce rRNA in total RNA samples. Libraries were prepared following the standard manufacturer's protocol. Library quality was evaluated using Qubit and TapeStation. Constructed libraries were sequenced on the NovaSeq 6000 Illumina platform, producing 2 × 100 nt stranded paired-end reads.

Raw total RNA-sequencing FASTQ files were aligned to the Macaca mulatta reference genome from Ensembl (Mmul_10, INSDC Assembly GCA_003339765.3) using STAR 2.7.8a in Partek, a bioinformatics software suite (Partek, Inc.). All samples had a quality score greater than 35, so no bases were trimmed, and 100% of sequencing bases overlapped with read lengths of the reference genome. Genes were filtered out of the count matrix if the sum of reads across all samples was less than 10. Downstream analysis of total RNA-sequencing count data was completed with the R statistical computing environment (R version 4.4.0). Gene counts were filtered by a row mean of 10 or greater, then normalized using the edgeR package to implement Trimmed Mean of M-values (TMM) normalization for technical biases. The voom function log-transforms the data and models the mean-variance relationship, which stabilizes the variance across different levels of gene expression. Differentially expressed genes (DEG) were identified between GBS infection groups and saline controls. Comparisons were based on a linear model fit for each gene using limma. P-values were obtained using limma's moderated t-test with empirical Bayes variance shrinkage. Genes were also filtered using the filterByExpr function in edgeR, and unannotated and mitochondrial or ribosomal genes were excluded when making the volcano plots. Significance required |fold change| ≥1.5 and adjusted $p<0.05$ (Benjamini-Hochberg). Differential expression counts reported in the text used a fold change of 1.5, while volcano plots (Fig. S5-S11) display a more stringent fold change of 2, after excluding unannotated, ribosomal, and mitochondrial genes.

**HDAC10 *Gene Expression qPCR on Chorioamniotic Membranes***

Gene expression by RT-qPCR was used to quantify the concentration of *HDAC10* in chorioamniotic membrane samples to validate total RNA-sequencing results. cDNA was synthesized utilizing 500ng of RNA isolated from homogenized chorioamniotic membrane samples. TaqMan Gene Expression Assay for *HDAC10* (Thermo Fisher Scientific, cat #4351372, assay ID: Rh02848946_m1) and *GAPDH* (Thermo Fisher Scientific, cat #4351372, assay ID: Rh02621745_g1) were used for primers and probe mix for the qPCR. TaqMan Fast Advanced Master Mix (Fisher Scientific, cat #444964) was used for amplification. qPCR was performed on the QuantStudio 3 machine (Thermo Fisher Scientific, cat #A28132) according to the manufacturer's instructions. Samples were run in triplicate. Positive results were defined as any sample with an average Ct value of 38 cycles or less. The log2 fold change was calculated using the $2^{-\Delta\Delta Ct}$ method. ΔCt was calculated for each group by subtracting the Ct value for *GAPDH* from the Ct value for *HDAC10*. The ΔΔCt was calculated by subtracting the ΔCt of the control group from the ΔCt of each GBS-infected group (localized, resolved, progressing). Then, the log2 fold change was calculated from the value of $2^{-\Delta\Delta Ct}$.

***Nanostring nCounter and Analysis***

nCounter was employed to validate and investigate immune changes at the maternal-fetal interface. A custom code set of 30 cell death genes and the Nanostring nCounter NHP Immunology V2 panel was created for analysis of total RNA from chorioamniotic membrane samples. Total RNA extracted from the myometrium samples was analyzed with the original Nanostring nCounter NHP Immunology V2 gene expression code set (Nanostring, cat #11500276). RNA hybridization reactions were completed based on the manufacturer's protocol. Hybridized probe and RNA complexes were immobilized on nCounter Cartridges using the nCounter Prep Station and quantified with the nCounter Digital Analyzer (Nanostring, cat #MAN-C0035-07).

Downstream analysis of raw nCounter data was completed in nSolver software (4.0). Background thresholding correction was applied based on negative controls, where probe counts below one were set to a value of 1. Normalization was completed using positive controls for technical variation and housekeeping genes for biological variation. The normalized count matrix was log2-transformed in the R statistical computing environment via the voom function. Comparisons were based on a linear model fit for each gene using the limma package. P-values were obtained using limma's moderated t-test with empirical Bayes variance shrinkage. Significance required |fold change| ≥1.5 and adjusted $p<0.05$ (Benjamini-Hochberg).

***Nanostring GeoMx Digital Spatial Profiling Whole Transcriptome Analysis***

Transcriptional profiles of the fetal and maternal placenta were delineated at spatial resolution. Formalin-fixed, paraffin-embedded chorioamniotic membranes and decidua tissue were sectioned onto microarray slides. The samples underwent deparaffinization and overnight in situ hybridization with UV-cleavable oligonucleotide probes based on the manufacturer's protocols. The microarray slides were incubated with morphological fluorescent antibodies used to highlight key histological features and draw the regions of interest (ROIs): amnion, chorion, and decidua. The morphological antibodies were (1) anti-pan cytokeratin-Alexa Fluor 488 (Pan-CK, clone AE1/AE3; Novusbio, cat #NBP2-33200AF488); (2) anti-fibroblast activation protein-Alexa Fluor 594 (FAP, clone SP325; Abcam, cat #ab311827); (3) DAPI for nuclei visualization (Thermo Fisher, Waltham, MA); and (4) anti-GBS-Alexa Fluor 647 (Abcam, cat #ab53584). After selection of ROIs on the GeoMx DSP instrument (Nanostring, cat #MAN-10152-01), UV light was illuminated and photocleaved DNA oligos were collected via aspiration.

Library preparation was completed based on the manufacturer's protocol (Nanostring, #MAN-10150-01). Constructed libraries were sequenced on the NovaSeq 6000 Illumina platform, producing 2 × 100 nt stranded paired-end reads. Raw FASTQ files were transformed to DCC files by DRAGEN via BaseSpace Sequence Hub. Downstream analysis was completed with the R statistical computing environment (R version 4.4.0). DCC, sample annotation, and PKC files were compiled to develop spatial objects for analysis with the NanoStringNCTools and GeoMxTools packages. The standR package was implemented for quality control, normalization, and batch correction with compatible subsequent linear modeling analysis (e.g., edgeR and limma-voom). Preliminary gene-level quality control filtered out genes found in less than 10% of ROIs with a minimal count of 1.5. ROIs were not filtered based on area or nuclear count, as the library size to cell count distribution followed a relatively smooth pattern. ROIs with a library size less than 20,000 were filtered. Gene counts were further filtered by a row mean of 10 or greater, then normalized using edgeR (TMM normalization). The voom function log-transforms the data and models the mean-variance relationship. DEG analysis was carried out between the progressing and contained infection groups. Comparisons were based on a linear model fit for each gene using limma; P-values were obtained using limma's moderated t-test with empirical Bayes variance shrinkage. Genes were filtered using the filterByExpr function in edgeR, and unannotated and mitochondrial or ribosomal genes were excluded when making the volcano plots. No genes were significant using the Benjamini-Hochberg method for calculating false discovery rate. Therefore, this analysis reported raw p values for the single gene analysis, and focused on results from the GSEA, which was adjusted.

***Statistical Analysis***

Differential gene expression across all platforms (total RNA-seq, nCounter, and GeoMx digital spatial profiling) was analyzed using linear models fit with the limma package. P-values were obtained from limma's moderated t-test with empirical Bayes variance shrinkage, and significance for differential expression required an absolute fold change of 1.5 and an adjusted p-value <0.05. GSEA (clusterProfiler, Hallmark and GO:BP) used false discovery rate (FDR) correction with adjusted p-value <0.05. Cytokine and chemokine concentrations were compared between groups using the tests indicated in the corresponding figure legends. Statistical analyses were performed in the R statistical computing environment (version 4.4.0).

**Table S1.** GBS SIP RT-qPCR Primers and Probes

| **Name** | **Sequence (5′→3′)** |
|---|---|
| GBS SIP Forward | GTTCCAGCAGCTAAAGAGGAAG |
| GBS SIP Reverse | CCGGTGCTACTTTAGCTACTGG |
| GBS SIP Probe | 5′-FAM-CACCAGCTTCTGTTGCCGCTGAAACACCAGC-BHQ1-3′ |

This table shows the sequence information used to generate the forward primer, reverse primer, and probes to detect the GBS SIP gene by RT-qPCR.

**Table S2.** Demographics and Experimental Characteristics

| Group | Group ID | Strain | Dam Age at Inoculation (years) | Gestational Age at Inoculation (days) | Interval Between Inoculation and Delivery (days) | Fetus | |
|---|---|---|---|---|---|---|---|
| | | | | | | Sex | Weight (g) |
| Control | CTL1 | N/A | 5.7 | 127.3 | 1.10 | M | 242.0 |
| Control | CTL2 | N/A | 6.0 | 133.4 | 1.00 | F | 264.0 |
| Control | CTL3 | N/A | 14.5 | 135.3 | 1.04 | F | 299.5 |
| Resolved | RES1 | GBSΔ*covR*Δ*cylE* | 13.7 | 132.4 | 1.04 | M | 267.0 |
| Resolved | RES2 | GBSΔ*covR*Δ*cylE* | 14.8 | 129.4 | 3.04 | F | 299.0 |
| Resolved | RES3 | GBSΔ*covR* | 5.2 | 132.4 | 3.25 | M | 366.0 |
| Resolved | RES4 | GBSΔ*covR* | 4.7 | 128.4 | 1.17 | M | 236.0 |
| Resolved | RES5 | GBSΔ*covR* | 11.6 | 129.3 | 1.05 | M | 200.0 |
| Localized | LOC1 | GBSΔ*covR* | 11.7 | 130.4 | 1.06 | F | 310.0 |
| Localized | LOC2 | GBSΔ*covR* | 7.0 | 130.3 | 1.05 | M | 310.0 |
| Localized | LOC3 | GBSΔ*covR* | 4.6 | 135.4 | 1.00 | M | 351.5 |
| Localized | LOC4 | GBSΔ*covR* | 4.4 | 132.4 | 1.00 | M | 344.8 |
| Localized | LOC5 | GBSΔ*covR*Δ*cylE* | 8.4 | 135.4 | 3.09 | M | 340.0 |
| Localized | LOC6 | GBSΔ*covR*Δ*cylE* | 8.3 | 129.4 | 3.04 | M | N/S |
| Progressing | PRO1 | GBSΔ*covR* | 5.5 | 130.4 | 0.23 | M | 299.0 |
| Progressing | PRO2 | GBSΔ*covR* | 5.9 | 134.4 | 1.21 | F | 263.0 |
| Progressing | PRO3 | GBSΔ*covR* | 7.4 | 127.4 | 1.00 | F | 255.7 |
| Progressing | PRO4 | GBSΔ*covR*Δ*cylE* | 8.7 | 131.4 | 3.04 | F | 299.0 |
| Progressing | PRO5 | GBSΔ*covR* | 6.2 | 129.4 | 2.03 | F | 197.0 |

This table shows maternal age at inoculation, gestational age at inoculation, and the interval between inoculation and delivery for each animal. Animals are grouped according to outcome classification (Control, Resolved, Localized, and Progressing) and identified by individual group IDs. The strain each animal was inoculated with is listed for infected animals (N/A: control animals were not inoculated with a pathogen). Fetal sex and weight at delivery are provided for each corresponding fetus. One fetal weight measurement was not available and is denoted as not sampled (N/S). Abbreviations: CTL, control; RES, resolved; LOC, localized; PRO, progressing; N/S, not sampled; M, male; F, female.

**Table S3.** Method of GBS Detection and Preterm Labor Status in the Control and Infection Groups

| Group | Group ID | Preterm Labor | GBS Detection by Swab of the Choriodecidual Inoculation Site | GBS Detected in AF by Culture (CFU/mL) | GBS SIP DNA in AF by qPCR (GE/mL) | GBS Detected in Fetus by Lung Culture (CFU/mL) |
|---|---|---|---|---|---|---|
| Control | CTL1 | No | Negative | 0 | 0 | 0 |
| Control | CTL2 | No | Negative | 0 | 0 | 0 |
| Control | CTL3 | No | Negative | 0 | 0 | 0 |
| Resolved | RES1 | Yes | Negative | 0 | 0 | 0 |
| Resolved | RES2 | Yes | Negative | 0 | 0 | 0 |
| Resolved | RES3 | Yes | Negative | 0 | 0 | 0 |
| Resolved | RES4 | Yes | Negative | 0 | 0 | 0 |
| Resolved | RES5 | No | Negative | 0 | 0 | 0 |
| Localized | LOC1 | Yes | Positive | 0 | 1.5E+002 | 0 |
| Localized | LOC2 | Yes | Positive | 0 | 3.3E+002 | 0 |
| Localized | LOC3 | Yes | Positive | 0 | 6.0E+002 | 0 |
| Localized | LOC4 | No | Positive | 0 | 7.0E+002 | 0 |
| Localized | LOC5 | No | N/S | 0 | 7.9E+002 | 0 |
| Localized | LOC6 | No | Negative | 0 | 1.1E+003 | 0 |
| Progressing | PRO1 | Yes | Positive | 2.90E+07 | 1.5E+006 | 2.20E+05 |
| Progressing | PRO2 | Yes | Positive | 9.40E+07 | 1.8E+006 | 9.90E+04 |
| Progressing | PRO3 | No | Positive | 7.90E+05 | 2.5E+006 | 4.80E+03 |
| Progressing | PRO4 | No | Positive | 1.00E+07 | 6.5E+007 | 9.88E+04 |
| Progressing | PRO5 | No | Positive | 9.20E+06 | 4.1E+007 | 3.60E+04 |

This table shows features of a progressing, localized, and resolved GBS infection based on a swab of the inoculation site, culture of the AF, GBS quantitative PCR (qPCR) on AF, and culture of fetal lung tissue. It also shows the occurrence of associated preterm labor per animal. Animals are grouped according to outcome classification (Control, Resolved, Localized, and Progressing) and identified by individual group IDs. One swab of the choriodecidual inoculation site was not available and is denoted as not sampled (N/S). Abbreviations: CTL, control; RES, resolved; LOC, localized; PRO, progressing.

**Figure S1.** Cytokines in Amniotic Fluid.

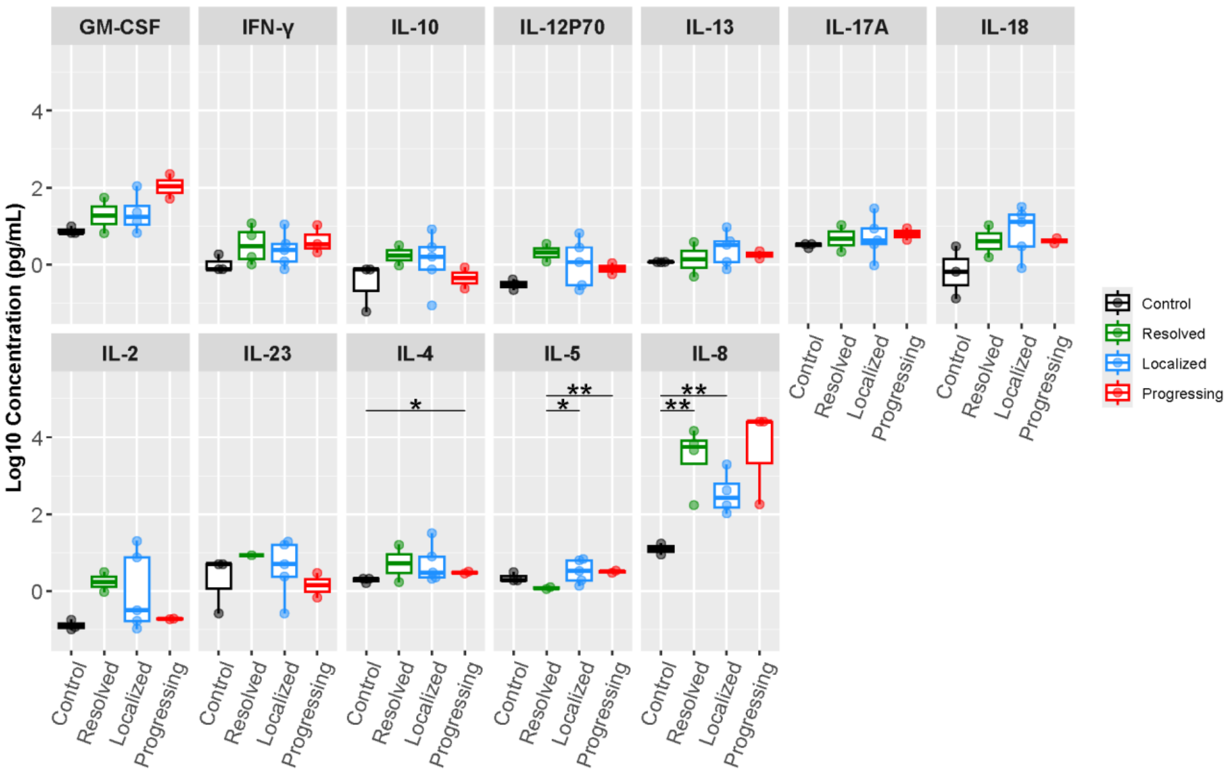


The box plots show cytokine concentrations from a 14-plex Luminex panel and an ELISA for IFN-β performed on amniotic fluid at the study endpoint or time of preterm labor. The x-axis shows the experimental groups, and the colors indicate uninfected controls (black), resolved (green), localized (blue), and progressing infections (red). The y-axis is the log10 of the cytokine concentration (pg/mL) with significant comparisons (unadjusted) shown using Student's t-test. A single star reflects a p-value <0.05 and double stars a p-value<0.01.

**Figure S2.** Cytokines in the Placental Chorioamniotic Membranes.

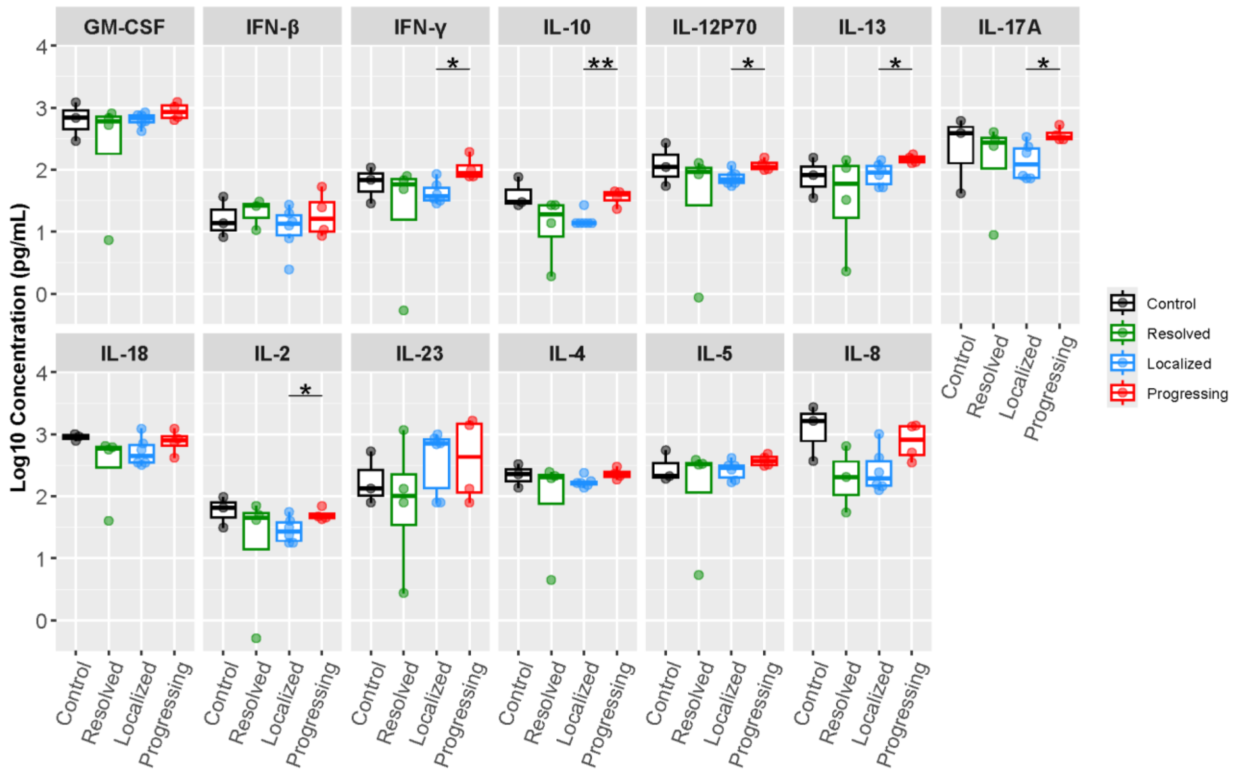


The box plots show cytokine concentrations from a 14-plex Luminex panel and an ELISA for IFN-β performed on homogenized tissues from the placental chorioamniotic membranes at the study endpoint or time of preterm labor. The x-axis shows the experimental groups, and the colors indicate uninfected controls (black), resolved (green), localized (blue), and progressing infections (red). The y-axis is the log10 of the cytokine concentration (pg/mL) with significant comparisons (unadjusted) shown using Student's t-test. A single star reflects a p-value <0.05 and double stars a p-value<0.01.

**Figure S3.** Cytokine Concentrations in the Lower Uterine Segment Myometrium.

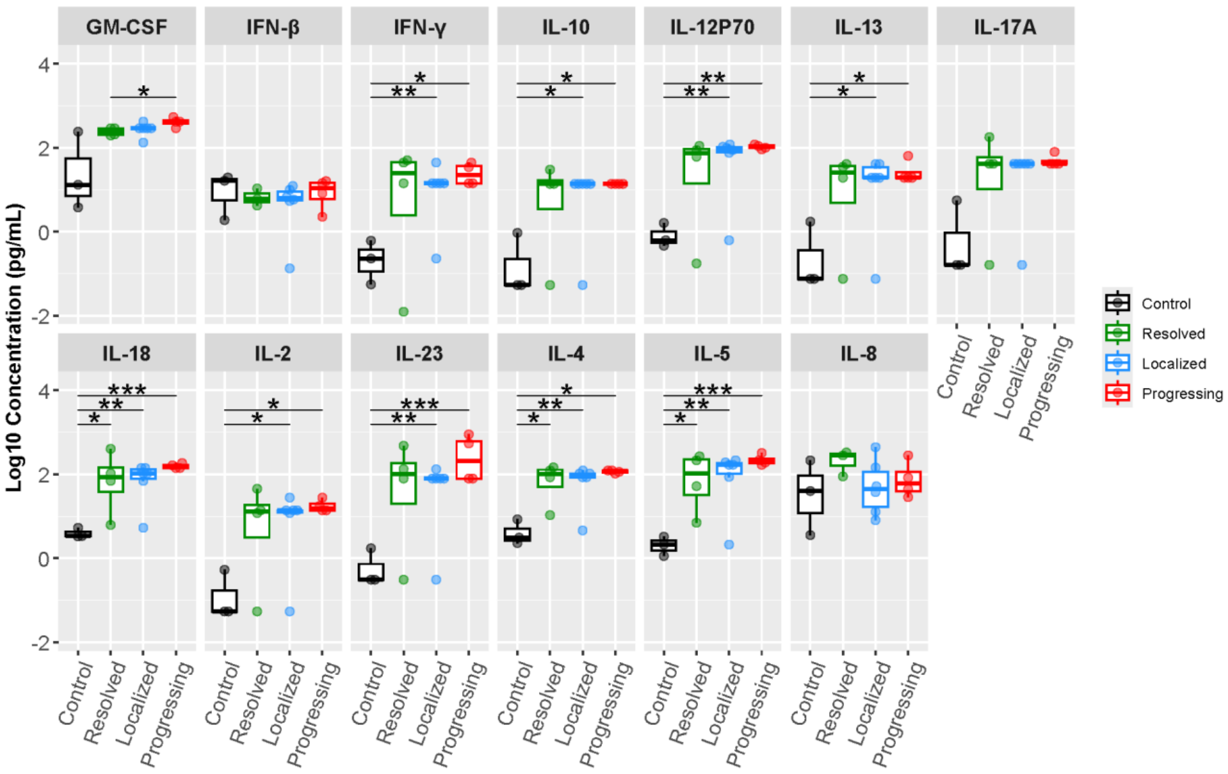


The box plots show cytokine concentrations from a 14-plex Luminex panel and an ELISA for IFN-β performed on homogenized tissues from the lower uterine segment myometrium at the study endpoint or time of preterm labor. The x-axis shows the experimental groups, and the colors indicate uninfected controls (black), resolved (green), localized (blue), and progressing infections (red). The y-axis is the log10 of the cytokine concentration (pg/mL) with significant comparisons (unadjusted) shown using Student's t-test. A single star reflects a p-value <0.05 and double stars a p-value<0.01.

**Figure S4.** PCA of Total-RNA-Seq from the Placental Chorioamniotic Membranes

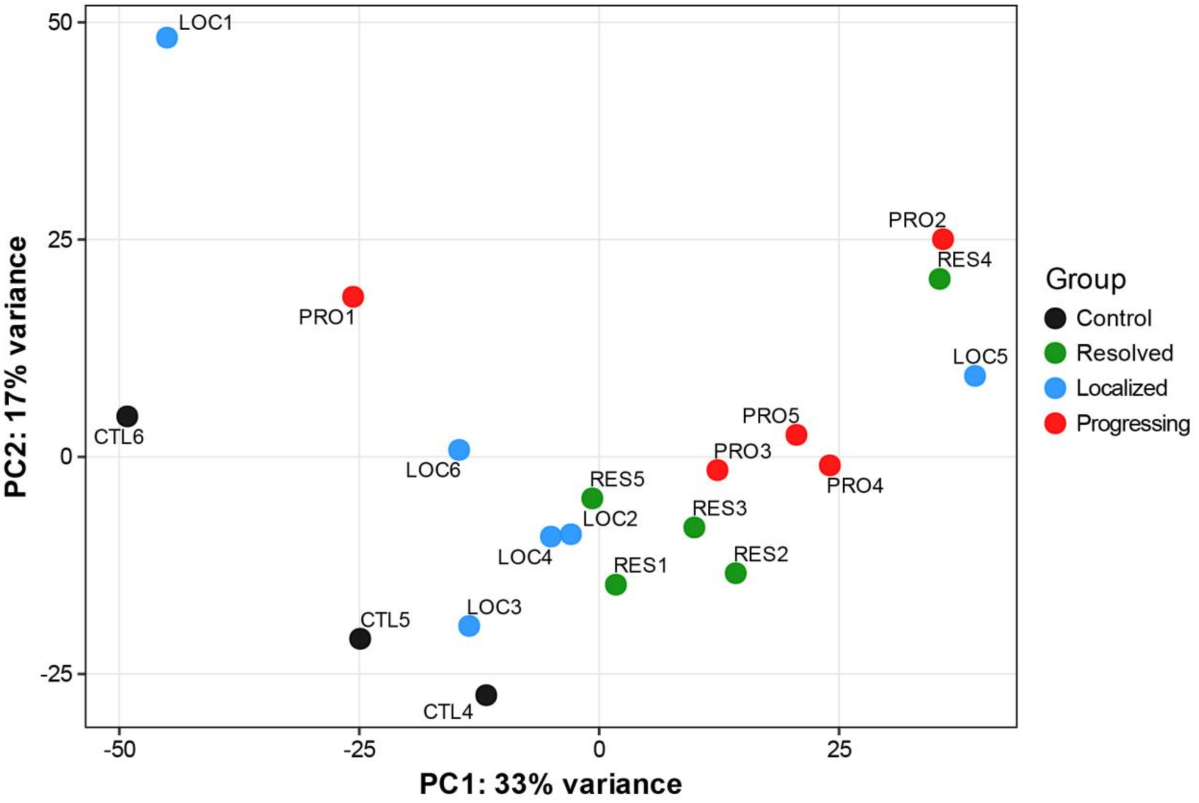


A principal component analysis (PCA) was performed on total RNA-Seq data from chorioamniotic membranes using normalized expression counts. Each point represents one animal and is colored by treatment group (black, control; green, resolved; blue, localized; red, progressing). The percent variance explained by each component is shown on the axis.

**Figure S5.** Total RNA-Seq Analysis of Progressing GBS Infections versus Control in the Placental Chorioamniotic Membranes at the Inoculation Site

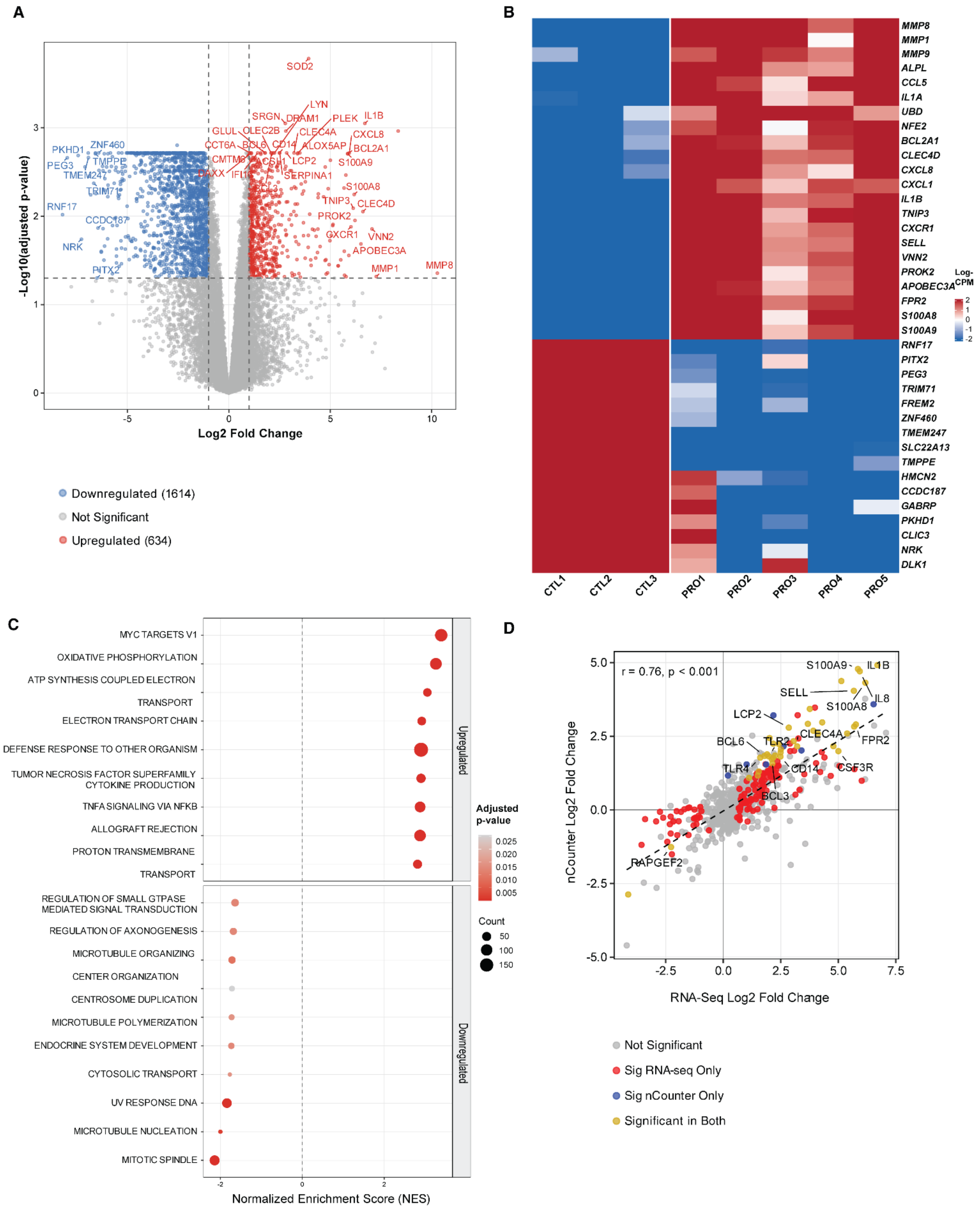

**A**: Volcano plot of significant differentially expressed genes found via total RNA-sequencing in Progressing vs Control animals within the placental chorioamniotic membranes at the inoculation site (>|2.0 fold change|, adjusted $p<0.05$). Genes that were not annotated, ribosomal, and mitochondrial genes were not shown or counted in the plot. The red color represents upregulated genes while blue corresponds to downregulated genes. **B**: Heatmap of top up- (red) and down-regulated (blue) genes showing normalized counts per animal. **C**: Top up- and down-regulated pathways as found via GSEA sorted by normalized enrichment score. Size of dot corresponds to number of genes in that pathway found to be enriched and the color denotes the adjusted p-value. **D**: Correlation plot showing a tight correlation between RNA-Seq and nCounter gene expression. Differentially expressed genes are shown as red if significant in RNA-Seq data only, blue if significant in nCounter data only, and yellow if significant in both data sets.

**Figure S6.** Total RNA-Seq Analysis of Localized GBS Infections versus Control in the Chorioamniotic Membranes at the Inoculation Site

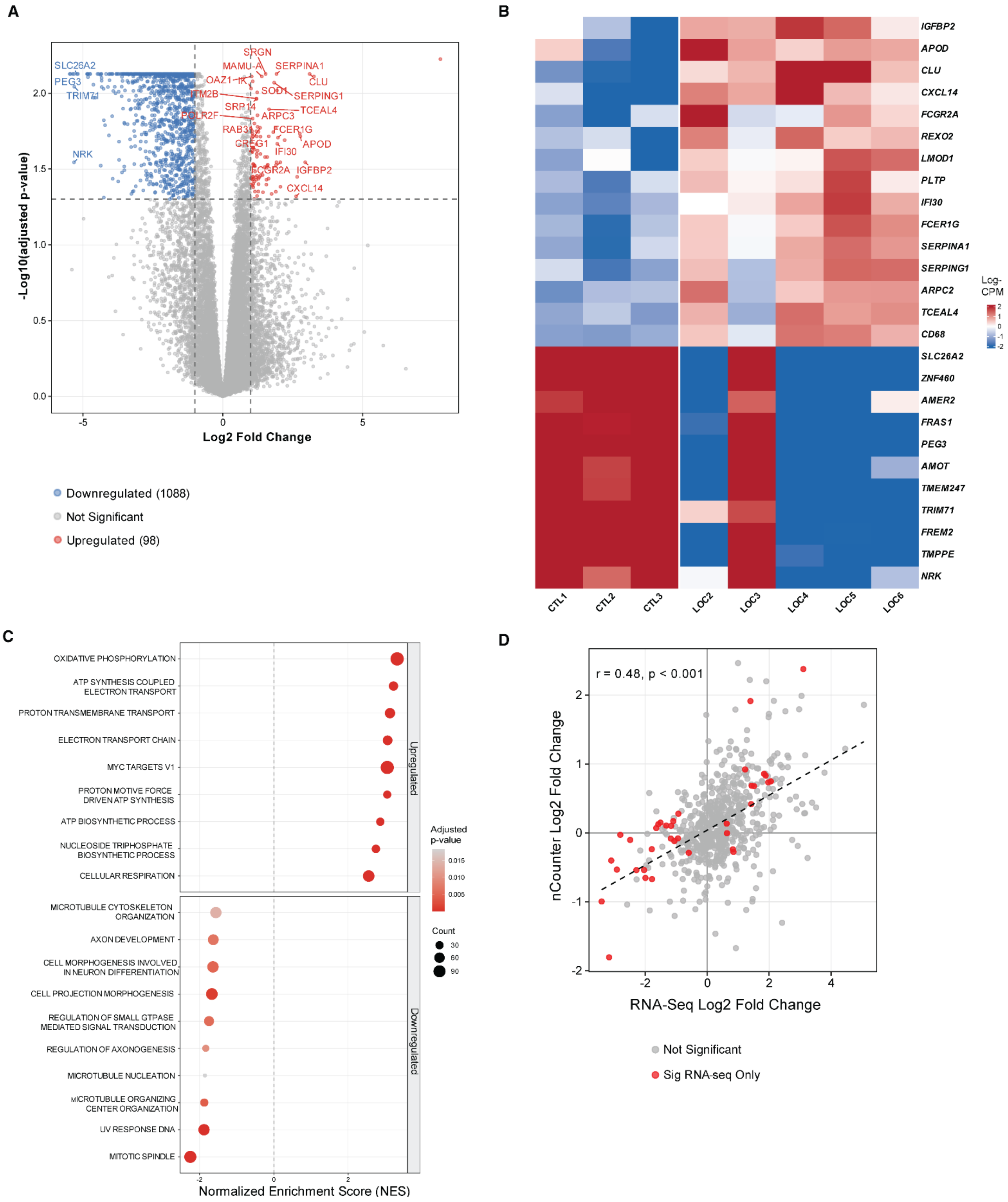

A: Volcano plot of significant differentially expressed genes found via total RNA-sequencing in Localized vs Control animals within the placental chorioamniotic membranes at the inoculation site (>|2.0 fold change|, adjusted $p<0.05$). Genes that were not annotated, ribosomal, and mitochondrial genes were not shown or counted in the plot. Red are upregulated genes while blue corresponds to downregulated genes. B: Heatmap of top up- (red) and down-regulated (blue) genes showing normalized counts per animal. C: Top up- and down-regulated pathways as found via GSEA sorted by normalized enrichment score. Size of dot corresponds to number of genes in that pathway found to be enriched and the color denotes the adjusted p-value. D: Correlation plot showing a tight correlation between RNA-Seq and nCounter gene expression. Differentially expressed genes are shown as red if significant in RNA-Seq data only. No genes were differentially expressed in the nCounter data.

**Figure S7.** Total RNA-Seq Analysis of Resolved GBS Infections versus Control in the Placental Chorioamniotic Membranes at the Inoculation Site

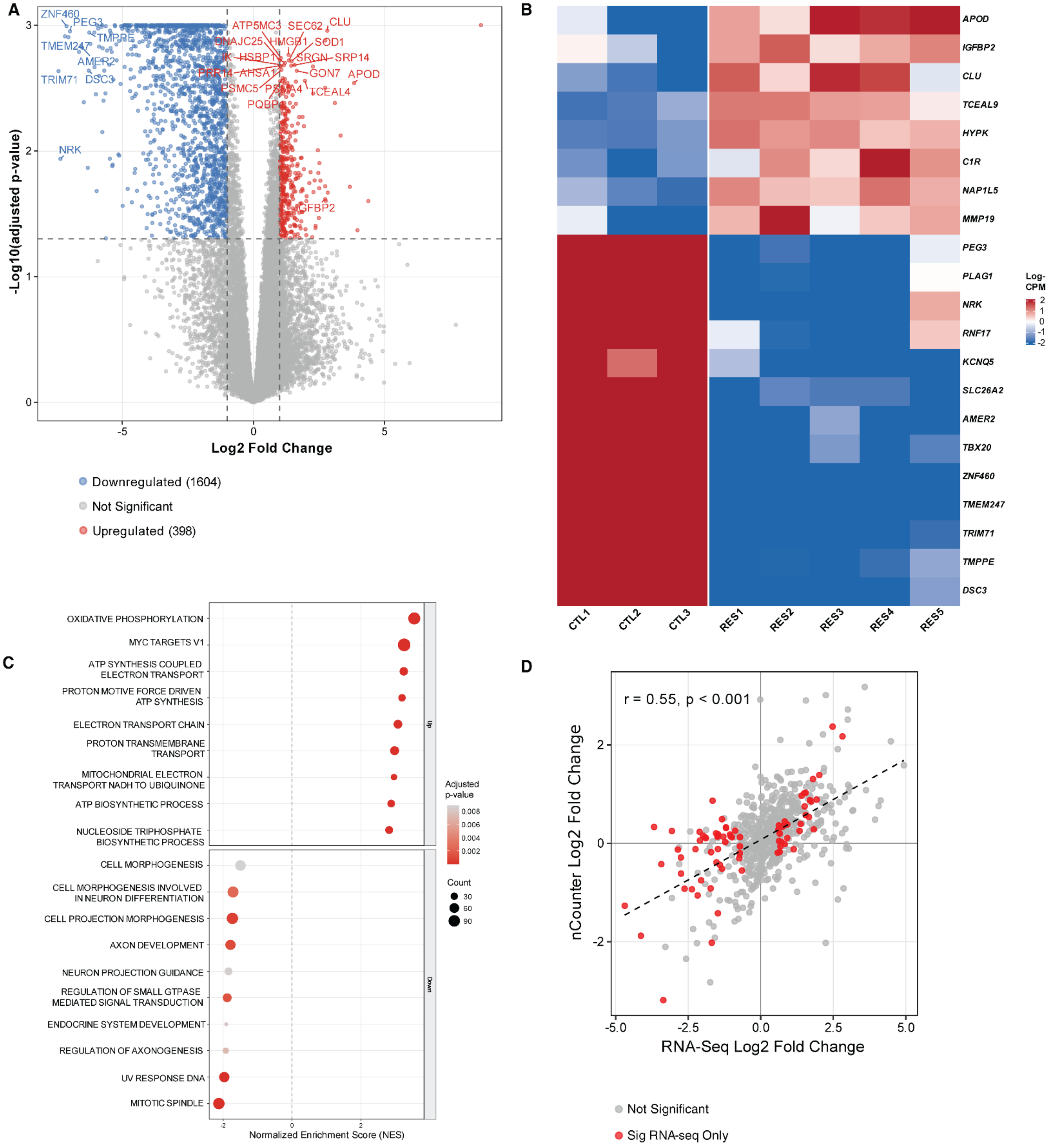

**A**: Volcano plot of significant differentially expressed genes found via total RNA-sequencing in Resolved vs Control animals within the placental chorioamniotic membranes at the inoculation site (>|2.0 fold change|, adjusted $p<0.05$). Genes that were not annotated, ribosomal, and mitochondrial genes were not shown or counted in the plot. The red color represents upregulated genes while blue corresponds to downregulated genes. **B**: Heatmap of top up- (red) and down-regulated (blue) genes showing normalized counts per animal. **C**: Top up- and down-regulated pathways as found via GSEA sorted by normalized enrichment score. Size of dot corresponds to number of genes in that pathway found to be enriched and the color denotes the adjusted p-value. **D**: Correlation plot showing a tight correlation between RNA-Seq and nCounter gene expression. Differentially expressed genes are shown as red if significant in RNA-Seq data only. No genes were differentially expressed in the nCounter data.

**Figure S8.** Total RNA-Seq Analysis of Progressing GBS Infections versus Resolved in the Placental Chorioamniotic Membranes at the Inoculation Site

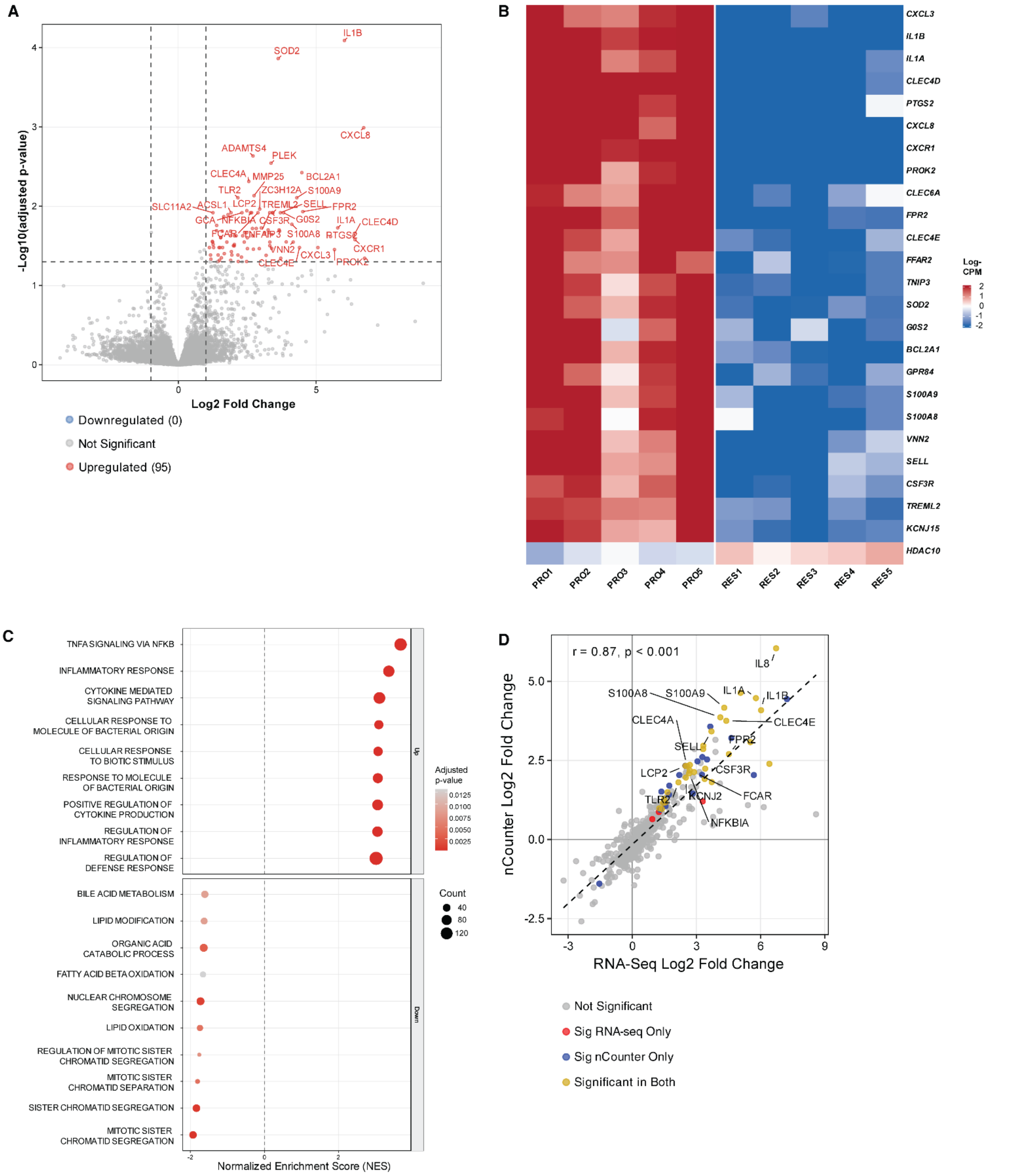

**A**: Volcano plot of significant differentially expressed genes found via total RNA-sequencing in Progressing vs Resolved animals within the placental chorioamniotic membranes at the inoculation site (>|2.0 fold change|, adjusted p<0.05). Genes that were not annotated, ribosomal, and mitochondrial genes were not shown or counted in the plot. Red are upregulated genes while blue corresponds to downregulated genes. Note that HDAC10 does not appear in the volcano plot because of the L2FC=-0.98 which was below the volcano plot significance cutoff of L2FC=1 (2-fold change). **B**: Heatmap of top up- (red) and down-regulated (blue) genes showing normalized counts per animal. **C**: Top up- and down-regulated pathways as found via GSEA sorted by normalized enrichment score. Size of dot corresponds to number of genes in that pathway found to be enriched and the color denotes the adjusted p-value. **D**: Correlation plot showing a tight correlation between RNA-Seq and nCounter gene expression. Differentially expressed genes are shown as red if significant in RNA-Seq data only, blue if significant in nCounter data only, and yellow if significant in both data sets.

**Figure S9.** Total RNA-Seq Analysis of Progressing GBS Infections versus Localized in the Chorioamniotic Membranes at the Inoculation Site

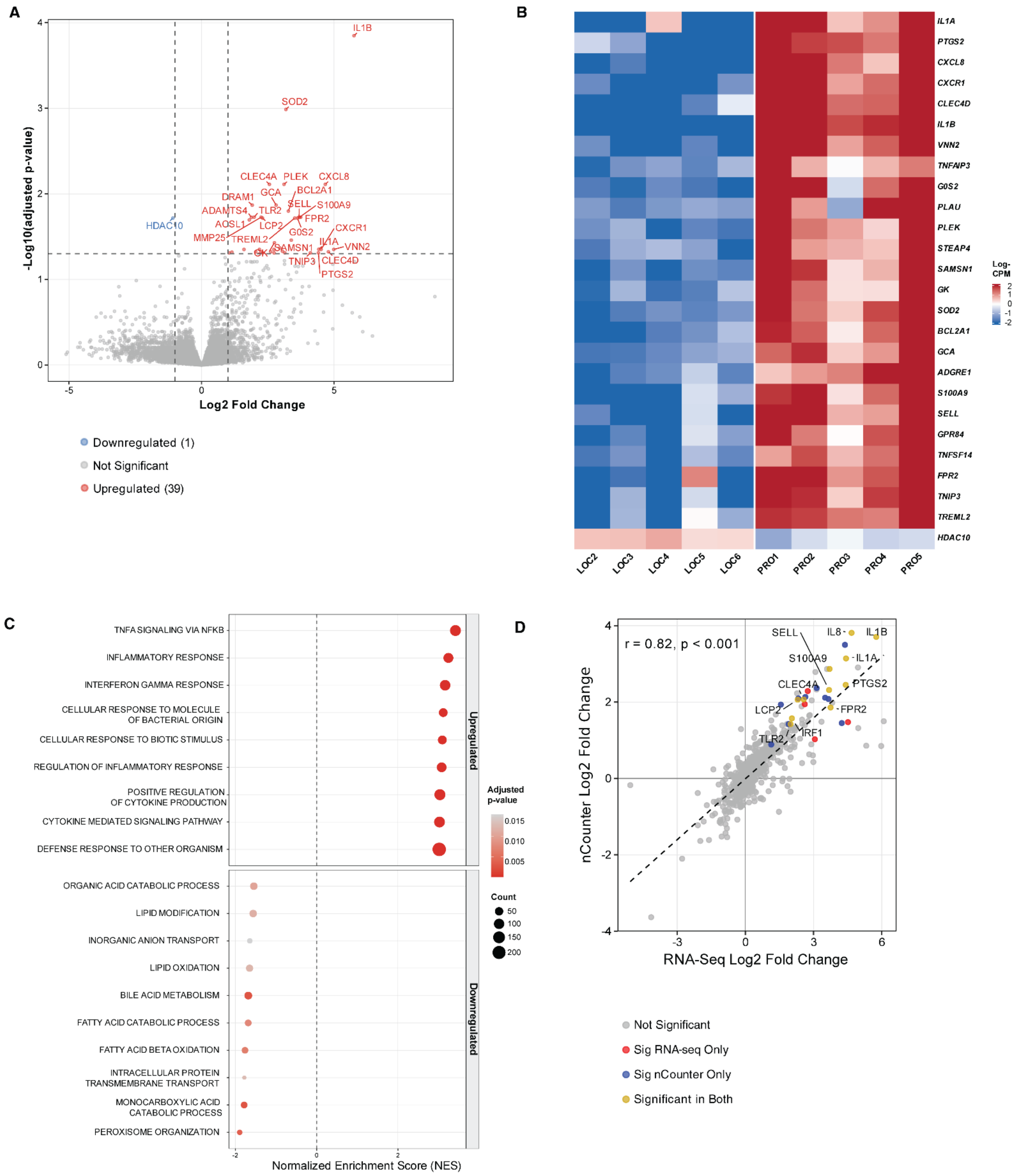

**A:** Volcano plot of significant differentially expressed genes found via total RNA-sequencing in animals with Progressing vs Localized GBS infections within the placental chorioamniotic membranes at the inoculation site (>|2.0 fold change|, adjusted p<0.05). Genes that were not annotated, ribosomal, and mitochondrial genes were not shown or counted in the plot. The red color represents upregulated genes while blue corresponds to downregulated genes. **B:** Heatmap of top up- (red) and down-regulated (blue) genes showing normalized counts per animal. **C:** Top up- and down-regulated pathways as found via GSEA sorted by normalized enrichment score. Size of dot corresponds to number of genes in that pathway found to be enriched and the color denotes the adjusted p-value. **D:** Correlation plot showing a tight correlation between RNA-Seq and nCounter gene expression. Differentially expressed genes are shown as red if significant in RNA-Seq data only, blue if significant in nCounter data only, and yellow if significant in both data sets.

**Figure S10**. Total RNA-Seq Analysis of Localized GBS Infections versus Resolved in the Chorioamniotic Membranes at the Inoculation Site

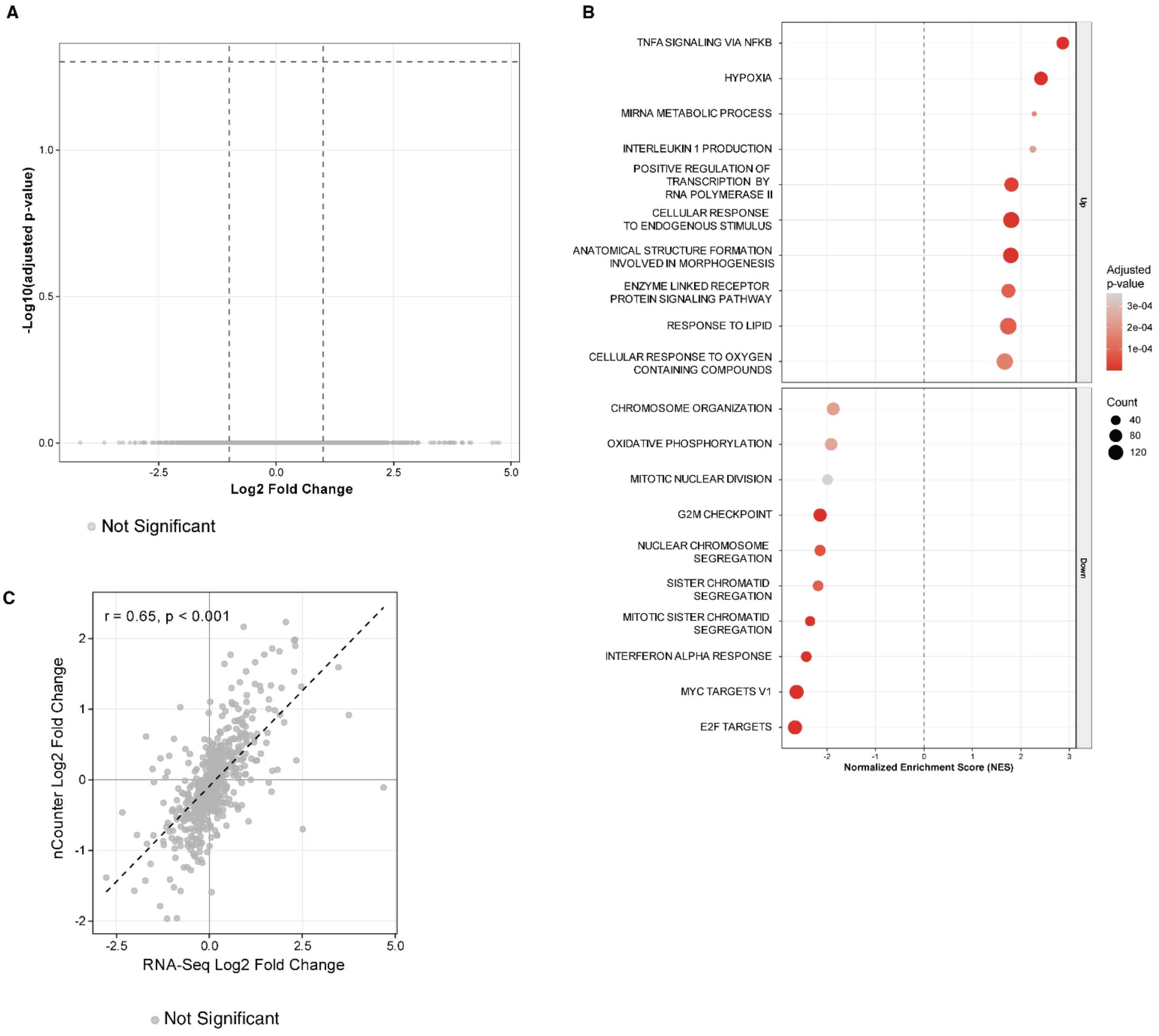


**A:** Volcano plot showing no differentially expressed genes in localized versus resolved infections in the chorioamniotic membranes. **B:** Top up- and down-regulated pathways as found via GSEA sorted by normalized enrichment score. Size of dot corresponds to number of genes in that pathway found to be enriched and the color denotes the adjusted p-value. **C:** Correlation plot showing a tight correlation between RNA-Seq and nCounter gene expression. All genes are colored gray, indicating that none were differentially expressed in either RNA-seq or nCounter.

**Figure S11.** Gene Expression of HDAC10 by RT-qPCR in Chorioamniotic Membranes at the Inoculation Site

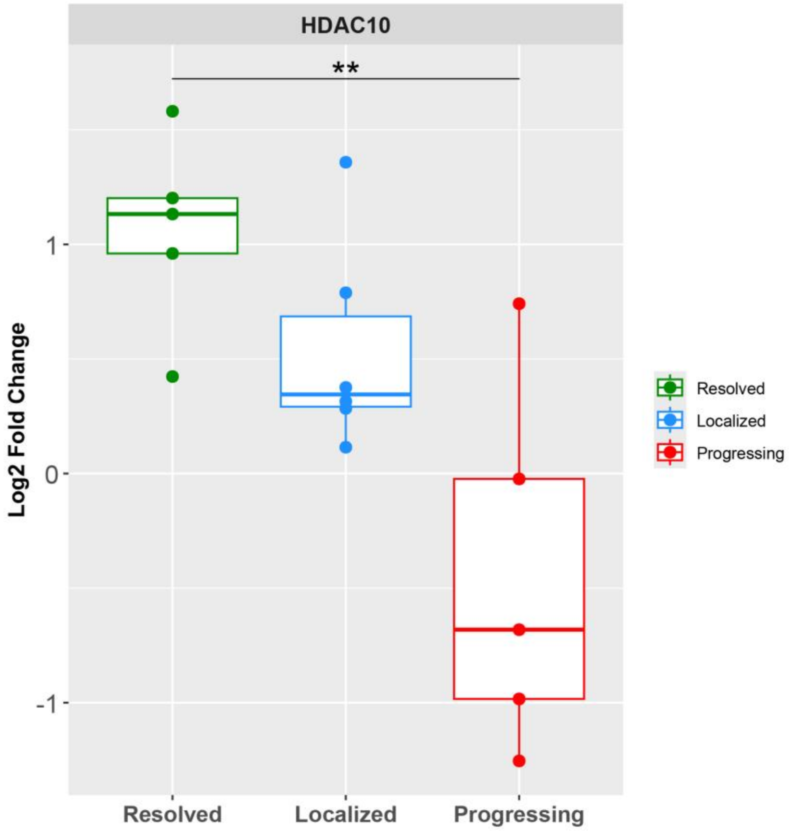


This boxplot shows the gene expression of HDAC10 assessed via RT-qPCR performed on homogenized tissues from the placental chorionic membranes at the study endpoint or time of preterm labor. The x-axis shows the experimental groups, and the colors indicate resolved (green), localized (blue), and progressing infections (red). The y-axis shows the log2 fold-change values of the ΔΔCt value comparing each sample to the control. Significance was calculated by Student’s t-test, with double stars reflecting a p-value<0.01.

**Figure S12.** Total RNA-seq Volcano Plots of GBS Infections and Uninfected Controls in the Myometrium Lower Uterine Segment Adjacent to the Inoculation Site

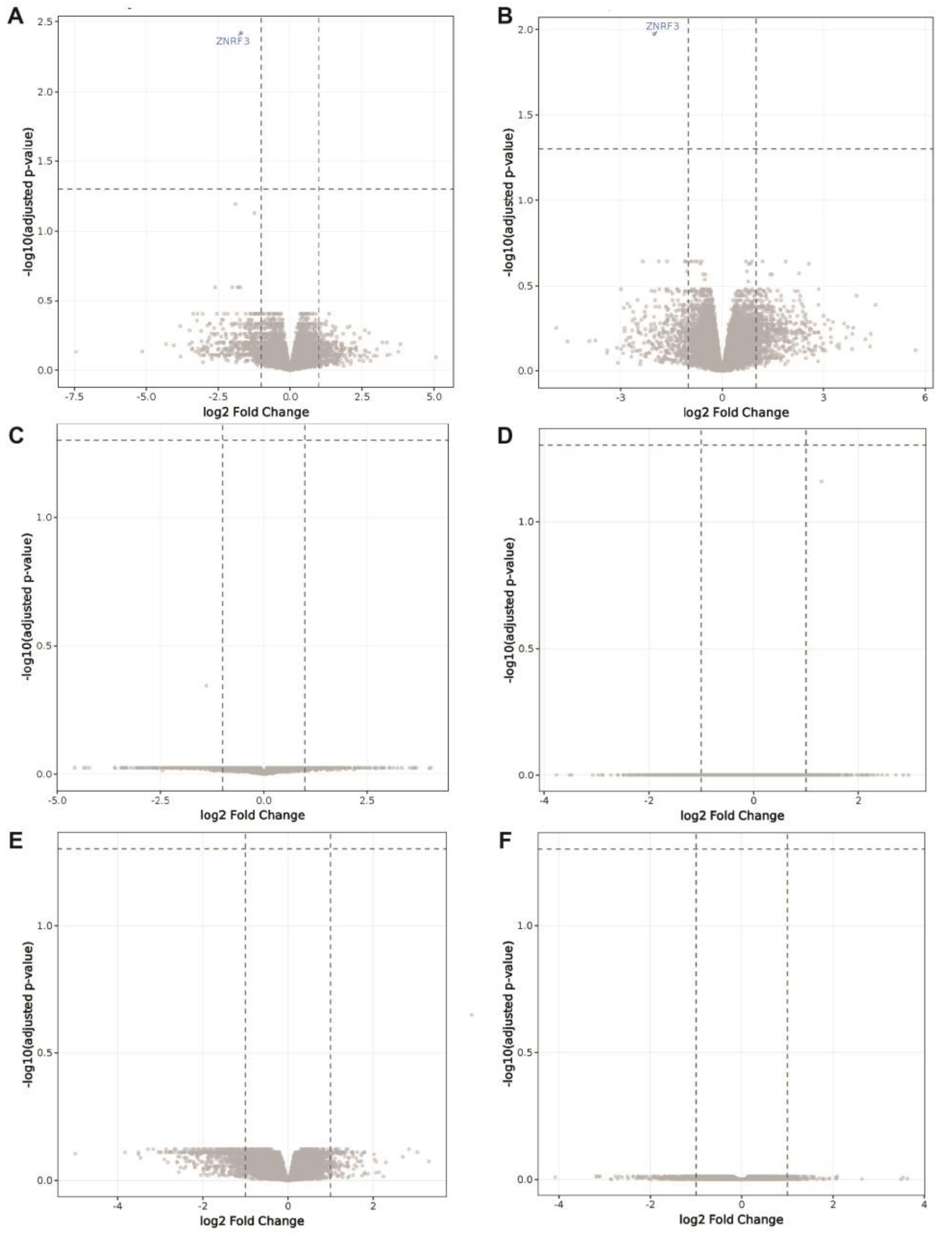


Volcano plots showing DEGs by contrast in the lower uterine segment myometrium: (A) localized versus control, (B) resolved versus control, (C) progressing versus control, (D) progressing versus localized, (E) progressing versus resolved, (F) localized versus resolved. No significant DEGs were identified in these contrasts with the exception of ZNRF3 downregulation in localized versus control (A) and resolved versus control (B).
